# Contrasting effects of glucose and methylglyoxal supplementation on blood oxidative status, blood cells’ telomere dynamics and apoptosis in birds

**DOI:** 10.64898/2026.07.09.737063

**Authors:** Adrián Moreno Borrallo, Roger Colominas-Ciuró, Bruno Colicchio, Radhia M’kacher, Anaïs Laurine Allak, François Criscuolo, Fabrice Bertile

**Author notes:** These authors share senior-authorship (François Criscuolo, Fabrice Bertile). Department of Ecology and Global Change, Desertification Research Centre -CIDE-, CSIC-UV-GVA, 46113, Moncada, Valencia, Spain.

## Abstract

Birds exhibit longer lifespans than similarly sized mammals, despite having higher mass-adjusted blood glucose levels. This makes them a valuable model for the comparative study of the metabolic and physiological aspects of aging. Circulating glucose contributes to multiple pathological processes, primarily through glycation reactions and the formation of advanced glycation end-products (AGEs), as well as by promoting oxidative stress. These mechanisms are interconnected by feedback loops and play a key role in the development of age-related pathologies. To explore the causal role of glycaemia in avian ageing, we conducted a one-year dietary supplementation experiment in captive zebra finches. Birds received either glucose- or methylglyoxal-enriched water. Previously, we observed that chronic glucose supplementation in zebra finches increased mortality, an effect that did not appear to be mediated by the associated increase in plasma protein glycation or AGE levels. Therefore, the mechanisms underlying increased mortality in the glucose group remained unclear. In the present study, we investigated how glucose and methylglyoxal supplementation affect blood oxidative status and red blood cell telomere dynamics and apoptosis. We found that methylglyoxal supplementation decreased the non-enzymatic antioxidant capacity (OXY) of plasma and increased DNA damage, while glucose supplementation had no significant effect on oxidative stress, although circulating glucose levels influenced oxidative status in a sex-dependent manner. Males exhibited a positive correlation between glucose levels and organic hydroperoxides and protein carbonyls. Additionally, we report, for the first time in birds, a seasonal variation in telomere length, which was more pronounced in glucose-supplemented individuals, yet seemed independent of oxidative status. Apoptosis probability increased with both treatments, particularly with the methylglyoxal supplementation. These results highlight that glucose and methylglyoxal trigger different glucotoxicity-related pathways, with distinct effects on bird health and aging. However, the relationship between glucose supplementation and mortality remains still unclear and warrants further investigation.

## Introduction

Birds are remarkable for their extended lifespans relative to their body mass compared to mammals (Lindstedt and Calder 1976), despite having physiological traits traditionally associated with reduced longevity (Munshi-South and Wilkinson 2010). For instance, birds exhibit a higher mass-specific basal metabolic rate than mammals (Lasiewski and Dawson 1967; Speakman 2005), a feature theoretically linked to higher oxidative damage production due to elevated oxygen consumption rates, according to the reactive oxygen species (ROS) theory of aging (Harman 1955). However, the relationship between metabolic rate and oxidative damage is complex, and the opposite relationship has been reported, i.e. higher metabolism related with lower oxidative damage (e.g. Colominas-Ciuró et al. 2022). Indeed, ROS production depends not only on oxygen consumption but also on how oxygen is utilized in metabolism, including factors such as mitochondrial uncoupling (Criscuolo et al. 2005a; Criscuolo et al. 2005b; Hou et al. 2021). Additionally, species-specific adaptations — such as membrane composition (Hulbert 2005; Hulbert et al. 2007), antioxidant defences, or damage repair efficiency (Hou and Amunugama 2015) can mitigate oxidative stress. Comparative studies have suggested that birds may experience lower oxidative stress, potentially due to reduced superoxide production or enhanced resistance to oxidative damage (reviewed in Skulachev 2004), although results vary across tissues and depending on the metrics used (Jimenez et al. 2019).

Another distinctive feature of birds is their elevated blood glucose levels, higher in average than in any other vertebrate group and double those of mammals (Polakof et al.2011). Glucose and other reducing sugars participate in non-enzymatic glycation reactions, forming adducts with compounds like proteins or nucleic acids (Maillard 1912; Suji and Sivakami 2004). These reactions can disrupt protein structure and function (e.g. Vetter and Indurthi 2011), and the situation can worsen due to oxidative stress, which promotes sugar autoxidation and the formation of reactive dicarbonyl compounds such as glyoxal and methylglyoxal. Reactive dicarbonyl compounds contribute to the formation of advanced glycation end products (AGEs; reviewed in e.g. Twarda-Clapa et al. 2022), which accumulate in tissues and are implicated in diseases such as atherosclerosis, kidney dysfunction and neuropathies (Chaudhuri et al. 2018), as observed in diabetes (Khalid et al. 2022).

Blood oxidative status is influenced by glycaemia in several animal species, including humans, mice and birds (e.g. Folmer et al. 2002; Menon et al. 2004; Vágási et al. 2020), and dicarbonyl compounds such as methylglyoxal are known inducers of oxidative stress and cytotoxicity (Chang and Wu 2006; Desai et al. 2010; Sena et al. 2012). Hyperglycaemia can further promote cell death through glycoxidative damage (Allen et al. 2005). Methylglyoxal has also been shown to induce eryptosis (erythrocyte apoptosis) in humans (Nicolay et al. 2006), with underlying mechanisms potentially mediated by superoxide production (e.g. Du et al. 2001). Elevated apoptosis levels may lead to anaemia and circulatory complications, such as increased thrombosis risk in diabetes patients (Williams et al. 2023). In birds, however, these pathological outcomes seem less likely to occur for a given glycaemia, despite their circulating glucose levels that would be considered pathological in humans. A recent comparative analysis suggests that blood glucose levels in birds are not associated with oxidative damage across species (Vágási et al. 2024), indicating potential adaptive mechanisms that mitigate the deleterious effects of circulating glucose. Identifying these adaptations is crucial to understanding how birds maintain high glucose levels while resisting associated damage.

To explore the effects of glucose and its derivatives on bird fitness and ageing, we conducted a year-long dietary supplementation experiment in captive zebra finches. Birds received either glucose or methylglyoxal through their drinking water. In a previous study (Moreno Borrallo et al. 2026), we demonstrated that glucose supplementation increased plasma protein glycation and mortality hazard, while both glucose and methylglyoxal elevated plasma AGE and affected physiological performance indicators, such as resting metabolic rate, flight speed and sexual signalling. Here, we investigated how these dietary supplements influence oxidative status and some of its expected consequences. Thus, we aim to determine whether oxidative status, including mitochondrial superoxide production, impacts telomere dynamics (Metcalfe and Olsson 2021), as it is often influenced by oxidative stress (Zglinicki 2002; Reichert and Stier 2017; Armstrong and Boonekamp 2023). Telomeres, which protect linear chromosomes from the end-replication problem, generally shorten with age (Chakravarti et al. 2021), and are hypothesized to mediate physiological and life-history trade-offs (Tobler et al. 2022). Given the high guanine content of telomeres, levels of 8-OH-d-oxo-Guanosine (our DNA damage marker) are expected to correlate with telomere damage and shortening in particular (Kawanishi and Oikawa 2004). We also assess the effects of supplementation on apoptosis, as while hyperglycaemia has been linked to anaemia in birds (Minias 2014), avian erythrocytes’ low glucose dependence (Johnstone et al. 1998) may confer resistance to hyperglycaemia-induced apoptosis. Finally, we evaluate whether these parameters contribute to the observed mortality patterns.

We hypothesized that glucose supplementation should induce higher oxidative stress than methylglyoxal, as glucose affects multiple pathways beyond reactive dicarbonyl compound production (Brownlee 2001). Additionally, glucose uptake by avian red blood cells is likely limited due to their low dependence on glucose as a metabolic fuel. Plasma oxidative stress markers might therefore be more affected by glucose than cellular oxidative stress markers (e.g. DNA damage). The effects of glucose on metabolic rate (Moreno Borrallo et al. 2026) may also lead to consequences that differ from those associated with methylglyoxal supplementation alone.

## Material and methods

### Animal model and study design

A population of 90 zebra finches (*Taeniopygia guttata*), originating from diverse sources (**Table ESM1.1**), was randomly assigned to three treatment groups (n= 30 per group, 15 males and 15 females) using the R function **sample()**. Birds were housed in outdoors aviaries at the IPHC – DEPE (Strasbourg, France), with one aviary per treatment: control (still water), glucose (50 g/L in drinking water; Sigma Aldrich G8270) and methylglyoxal (8.33 g/L; dilution prepared from Sigma Aldrich M0252). The concentrations were determined during a pilot study (see Moreno Borrallo et al. 2026). The experiment was carried out from September 21^st^ 2022, to September 25^th^ 2023.

The age of 86 of the 90 birds was known, most of them belonging to two main age classes: young (∼4-year-old) and old (∼6-year-old) individuals. Age distribution was balanced across treatment groups (see **Table ESM1.2** for detailed ages). Every three months (i.e. August and November 2022 and February, May and August 2023), we measured ageing-related parameters. For further details on housing conditions and general experimental protocols, including blood sampling see Moreno Borrallo et al. 2026.

#### Sampling Schedule

Each sampling season lasted several weeks. The first 1-2 weeks were dedicated to ageing-related measurements (metabolic rate, beak coloration and flying performance), which are discussed elsewhere (see Moreno Borrallo et al. 2026). We nevertheless tested here the effects of some of these parameters on oxidative status (see **Statistics** section). This was followed by two weeks of blood sampling. All oxidative stress, apoptosis and telomeres measurements discussed in this manuscript were performed on blood collected during the second week of blood sampling. Due to technical constraints, cytometry measures (apoptosis and mitochondrial superoxide production) were not available at baseline (August 2022). Oxidative status parameters were assessed every six months, corresponding with baseline, midpoint and shortly before the end of the experiment (i.e. August 2022, February and August 2023). Telomere length analyses were performed on a subset of the initially sampled birds (n = 221 samples: Baseline = 76, November = 48, February = 43, May = 39, August = 15).

### Oxidative status measurements

#### Plasma oxidative status

In bird plasma, we assayed the non-enzymatic antioxidant capacity (OXY, Diacron®), levels of total organic hydroperoxides (d-ROMs test; Diacron®) (see e.g. Alberti et al. 2000; Costantini 2016) and protein carbonyl levels (MAK486 – Sigma Aldrich). Plasma measurements required 20 µL of plasma per bird (8 µL for OXY, 2 µL for d-ROMs and 10 µL for protein carbonyls, including duplicates). If an insufficient volume was available (i.e. less than 16 µL), OXY and d-ROMs were prioritized over protein carbonyl assay. For volumes between 20 and 16 µL, protein carbonyls were assessed, but the duplicate for OXY was omitted. The absorbance was measured using a Tecan Infinite M200® microplate reader. Protein carbonyl levels were normalized by protein concentration, assessed afterwards from 5 µL of sample diluted on 5 µL of distilled water, using BCA assay (bicinchoninic acid; Thermo Fisher Scientific: Pierce™ BCA Protein Assay Kit, Catalogue Numbers 23225 and 23227) and a BSA standard curve.

#### DNA damage

In bird red blood cells, we assayed cellular mitochondrial superoxide production (MitoSox^TM^ fluorescence) and DNA damage levels (8-d-Oxo Guanosine, EpigenTek® kits). For DNA damage, DNA was extracted from 5 µL aliquots of unfrozen blood cells using an EpiQuik® kit (EpigenTek), proteinase K and RNAse (Macherey-Nagel®). DNA quality was verified using a Nanodrop® (absorbance ratios 260/230 nm and 260/280 nm > 2). The extraction was repeated if DNA concentration was lower than 100 ng/µL. DNA solutions were stored at -80°C until analysis. Most measurements were performed in March/April 2023 and August 2023, except for August 2023 samples, analyzed in April 2024 due to organizational constraints. The absorbance was measured using a Tecan Infinite M200® microplate reader.

### Mitochondrial superoxide production and apoptosis using flow cytometry

For superoxide production and apoptosis assessment, 10 µL of heparinized blood were diluted with 390 µL of cell culture medium (Dulbecco Modified Eagle Medium (DMEM) high Glucose, L-Glutamine, Sodium Pyruvate, Dominique Dutcher®, Brumath, France) previously brought to room temperature. Then, 75µL of each individual sample were dispensed in duplicate into a 96-well plate, to which 2.5 μl of a solution of 0.5 mM MitoSOX™ Red reagent (for mitochondrial superoxide determination) were added. This MitoSOX™ solution consisted of 130 μl of DMSO (dimethylsulfoxide) added to one tube of MitoSox™ Red (50μg). The MitoSox™ stock solution was then distributed in 4 aliquots of 30 μl to be stored at -20°C for further use, and the plate with the samples was sealed and incubated 30 min at 40°C in the dark. After this first incubation step, the plate was centrifugated at 2000 g at room temperature, the medium was discarded and cells were washed using 75 µL PBS. This wash step was repeated once again.

The washed cells were then resuspended with Annexin V binding buffer (previously diluted 10x, 0.1 M Hepes pH 7.4, 1.4 M NaCl, 25mM CaCl_2_) containing annexin-V-FITC (5μl of the stock solution of Annexin V and 1 µg/mL of propidium iodide, BD Biosciences), for rate of apoptosis determination. Annexin V binds to phosphatidylserine that becomes exposed on the outer layer of the plasma membrane in apoptotic cells. Propidium iodide (PI) enters necrotic cells but is excluded from apoptotic cells. As such, using flow cytometry (Accuri^TM^ C6 flow cytometer, BD, Oxford, UK), Annexin V binding and Propidium iodide uptake and associated fluorescence signals allow to distinguish living (not stained), pre-apoptotic (Annexin V stained), apoptotic (both Annexin V and propidium iodide) and dead cells (propidium iodide).

After Annexin V and propidium iodide addition, the plate was then sealed and incubated at room temperature in the dark for 15 min. Cells were harvested following a centrifugation step at 2000 g at room temperature, washed using 75 µL PBS, re-centrifuged at 2000 g, and finally resuspended with cell culture medium (75 µL).

Reading of fluorescence signal was performed thereafter by the cytometer Accuri^TM^ C6 flow cytometer (Becton Dickinson, BD, Franklin Lakes, NJ, USA). Up to 20000 red blood cells (RBC) were counted for each measurement, and a rinse cycle was added after each sample reading to avoid inter-samples’ contamination. We used 488 nm excitation for Annexin V, propidium iodide and oxidized MitoSox™ Red. In each plate, control samples including negative (no staining), or cells stained only with MitoSox™, Annexin V and PI were run to perform signals’ compensation. Signal detection was done using: FL1-FITC 530 nm for Annexin, FL2-PE-A (580 nm) for MitoSox™ Red, and FL4-APC-A (625 nm) for propidium iodide. To optimize cytometry results, performance validation and fluidic maintenance (including peristaltic pumps) were done regularly prior to experimental sessions. Samples were analysed at low flow rate, using a threshold of 25000 to exclude debris and electronic noise. Cell doublets were excluded using a first gating on the FSC-A/FSC-H profiles. Analysis of the data was done using plot cell counts for Annexin V and propidium iodide to get living (no staining) and early apoptotic (Annexin V stained) cells proportions. For both cell categories, we determined the mean MitoSox™ intensity (mean values of superoxide production). Finally, cells were subsequently stored at 6 °C for the measurements to be repeated after 24h to get an estimate of the dynamic of apoptosis and mitochondrial superoxide production.

### Quantitative analysis of telomere length using Q-FISH

#### Cytogenetic sample preparation

The process followed was the same as in (M’kacher et al. 2020). Briefly, two drops of blood were added to RPMI1640 medium, centrifuged (7 min at 1400 rpm, room temperature), and the cell pellet was resuspended in 0.075 M potassium chloride (KCl) (Merck, New Jersey, US). After a 20-min incubation in a 37 °C water bath (hypotonic shock), cells were pre-fixed by adding approximately five drops of fixative (3:1 ethanol/acetic acid) to each tube under agitation, and the tubes centrifuged for 7 min at 1400 rpm at room temperature. The supernatant was then removed and cells were re-suspended in the fixative solution followed by another centrifugation under the same conditions. After two additional rounds of these fixation steps, the cells were stored in the fixative solution at 4 °C overnight and then spread on cold wet slides the next day. The slides were dried overnight at room temperature and stored at −20 °C until further use.

#### Telomere staining

Telomeres were stained with a Cy-3-labelled PNA probe specific for telomeres (Aging kit; Cell Environment, Evry, France), as previously described (M’kacher et al. 2020). Briefly, slides were washed with 1X PBS and fixed with 4% formaldehyde at room temperature. After rinsing three times with PBS, they were treated with pepsin (0.5 mg/ml) at 37°C for 5 min. After an additional rinsing three times with PBS, the slides were sequentially dehydrated with 50%, 70%, and 100% ethanol and air-dried. The telomere and centromere probes were added to the slides and subsequently denatured on a hot plate at 80°C for 3 min and then incubated in the dark for 1 h at room temperature. The slides were subsequently rinsed with 70% formamide/10 mM Tris pH 7.2 three times during 15 min and then in 50 mM Tris pH7.2/150 mM NaCl pH 7.5/0.05% Tween-20 (3 x 5min). After a final rinse with PBS, the slides were counterstained following with DAPI and mounted in PPD at the appropriate pH.

#### Telomere quantification

Quantitative image acquisition from slides with stained cells was performed using MetaCyte software (MetaSystem, version 3.9.1, Altlussheim, Germany). The exposure and gain settings remained constant between captures. The analysis of telomere signals was performed using TeloScore Software (Cell Environment, Paris France). The mean fluorescence intensity (FI) of telomeres was automatically quantified and analyzed in 10000 nuclei on each slide. The measurements were performed on triplicate slides.

### Statistical analyses

All statistical analyses were conducted using R (v4.5.1) with RStudio. General linear mixed models were fitted using **lmer()** function from **lme** package (Bates et al. 2015), unless otherwise specified. For each dependent variable, OXY, d-ROMs, Protein carbonyl, DNA damage, telomere length, apoptosis and mitochondrial superoxide production in both living and early-apoptotic cells, two distinct models were constructed.

The first model included treatment, month of measurement, sex and all their interactions as fixed effects, alongside a horizontal age component. The second model replaced the month with a longitudinal age component and its interaction with horizontal age, allowing for the estimation of potential senescence acceleration effects. Bird ID was consistently included as a random factor to account for repeated measures. Protein carbonyl, d-ROMs, and DNA damage levels were log_10_ transformed to meet model assumptions. OXY was included as a covariable in models predicting d-ROMs and protein carbonyl values, while DNA quantity (DAPI) was included as a log_10_ transformed covariate in models predicting telomere length. All covariables were centred to facilitate interpretation of the intercepts (Schielzeth 2010), except for longitudinal age, where zero was set at the start of the experiment.

Following the detection of a potential seasonal cycle in telomere length, a quadratic component of longitudinal age was incorporated into the complete model for telomeres, accounting for longitudinal age with both the linear and quadratic terms centred. For cytometry measures, time (0 vs. 24 hours) was included as an explanatory variable, along with its interactions. To account for the nested structure of the data, a random effect of month within individual (i.e. 1|Bird_ID/Month) was included in the models, reflecting the fact that each bird was measured twice per month.

For apoptosis, a betabinomial model with logit link was employed using the **glmmTMB()**package (Brooks et al. 2017; Kristensen et al. 2016). Both living and early-apoptotic cells were considered as dependent variables (using **cbind()**). The adequacy of these apoptosis models was assessed using the **testUniformity()** and **testDispersion()** functions from the **DHARMa** package (Hartig 2024).

For mitochondrial superoxide production, separate models were constructed for the living and the early apoptotic cells. These models included the number of cells, month and time (0 vs 24 hours) and their interactions as fixed effects. MitoSox signal values were log_10_ transformed. After observing a potential quadratic pattern, the square of the centred longitudinal age was also tested in the complete model accounting for longitudinal age for superoxide production in early apoptotic cells.

An additional set of models was developed to analyse the dynamics of superoxide production in both living and early-apoptotic cells (as in e.g. Lorenz et al. 2017; Cuong et al. 2007; Pearson et al. 2014; Francois Criscuolo et al. 2010). Residuals were extracted from models on MitoSox signal on the corresponding cell type both at zero and 24h, including only the interaction of the number of cells and month as a predictive covariate, and bird identity as a random intercept. This approach corrected for the difference in the relationship between MitoSox signal and cell number across months, as well as differences in cell number between different time points (0 vs 24 h), so that directly calculating the difference in MitoSox signal would not be appropriate. The difference in these residuals (after minus before), after correcting by regression to the mean (Verhulst et al. 2013; see telomere dynamics below), was then used as the dependent variable in models equal to the previously described ones, but now excluding the number of cells and time as explanatory variables.

Model selection was performed using the *dredge()* function from the *MuMIn* package (Burnham and Anderson 2002), with the final model chosen based on the lowest AICc value. Treatment, month, and cell number (for superoxide production models), along with their interactions, were retained in all models by principle. Marginal means and slopes were estimated using the *emmeans()* and *emtrends()* functions, and pairwise comparisons were conducted across treatments within month and across months within treatment. The significance was always set at α < 0.05. The final models selected, including all variables, are detailed in **ESM2.Box 1**.

#### Additional models

Additional models were constructed to explore the effects of DNA damage on telomere length, telomere dynamics (changes in telomere length) and apoptosis, as well as the effects of telomere length on apoptosis. For telomere dynamics models, adjustments were made for telomere length prior to the measured change, after correcting for regression to the mean effects with the equation from Verhulst et al. (2013), although in the opposing order (i.e. X_2_-X_1_), to test for possible increase in length. Furthermore, the effects of other variables measured in these birds (see Moreno Borrallo et al. 2026 for full explanation on such measures) were tested. These models included testing the effects of whole body oxygen consumption (VO_2_), measured on Resting Metabolic Rate (RMR) conditions, on blood cells mitochondrial superoxide production and oxidative damage variables (d-ROMs, protein carbonyls and 8-oxo-d-guanosine), whole blood glucose or plasma glucose (see Moreno Borrallo et al. 2026) on oxidative damage variables, and finally oxidative status variables (i.e. damage variables plus OXY) on albumin glycation rates and plasma Advanced Glycation End-products (AGE) levels, while adjusting by glucose levels (see Moreno Borrallo et al. 2026). Bird identity was included as a random intercept in all these models.

Finally, the effects of OXY, d-ROMs, protein carbonyl, DNA damage, and telomere length on survival were estimated using time-varying Cox proportional hazard models, with the **coxph()** function, controlling for OXY in the d-ROMs model and for age in all models, as described in Moreno Borrallo et al. (2026).

## Results

### Plasma oxidative status

#### OXY (Total Plasma Non-Enzymatic Antioxidant Capacity)

Total plasma non-enzymatic antioxidant capacity (in micromoles of HClO equivalents per mL) showed significantly lower levels at the end of the experiment (August 2023) in methylglyoxal treated birds compared with their previous levels (**Figure 1.A.,** Baseline: difference ± SE = 59.491 ± 14.5; t = 4.109, P = 0.0002; February: 62.746 ± 15.0; t = 4.188, P = 0.0002) and with levels in the control and glucose groups in August (Control: 46.009 ± 18.3; t = 2.516, P = 0.034; Glucose: 55.728 ± 21.8; t = 2.56, P = 0.03). These differences are estimated subtracting the August methylglyoxal group values from the other groups or time points.

**Figure 1.**
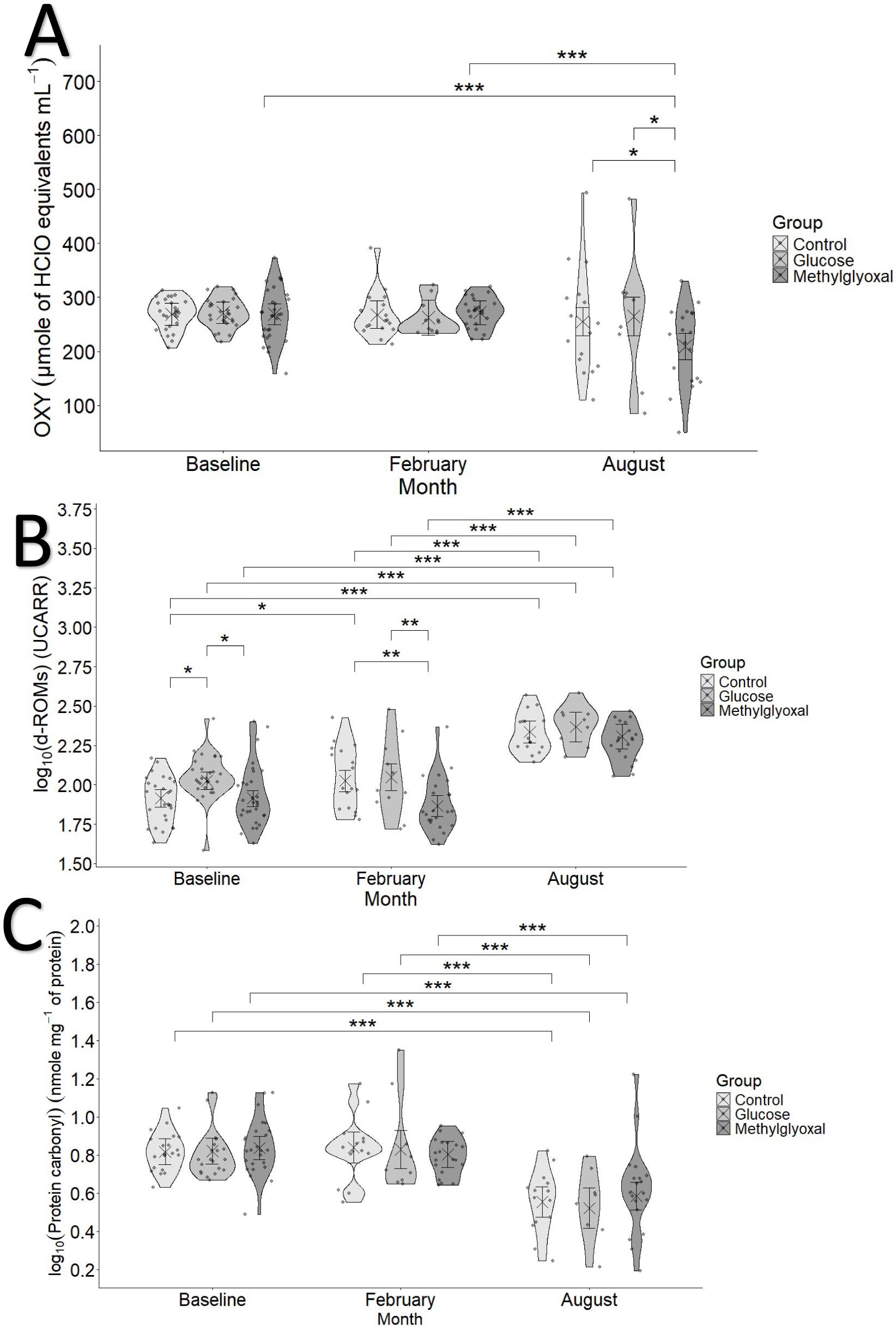
Plasma oxidative status markers along the experiment and across treatment groups. (**A**) OXY, (**B**) d-ROMs and (**C**) protein carbonyls. Crosses and error bars represent model-estimated marginal means ±95% CI. Significance annotations are based on pairwise contrasts performed separately within months and within treatments.

A longitudinal age effect was also observed: negative (i.e. a decrease along the experiment) for the methylglyoxal group (estimate ± SE: -77.1 ± 31.5; CI_95_[-139.6, -14.666]) and positive for the control group (106.3 ± 37.4; CI_95_[32.2, 180.378]), although these age effects were restricted to females. Repeatability was moderate but significant (R = 0.212, SE = 0.097, CI_95_[0.015, 0.397], P = 0.002).

#### d-ROMs (Total Plasma Organic Hydroperoxides)

The total plasma organic hydroperoxides significantly increased in all treatments at the end of the experiment compared with respective baseline and midpoint values, while control birds also showed increased d-ROMs in February compared with baseline (**Figure 1.B.** and **Table ESM2.1**). Additionally, methylglyoxal-treated birds exhibited significantly lower levels than birds in the control (Control - Methylglyoxal difference ± SE: 0.159 ± 0.048; t=3.297, P = 0.003) and glucose groups (Glucose - Methylglyoxal: 0.182 ± 0.054; t = 3.383, P = 0.002) at midpoint. Glucose treated birds showed higher d-ROM levels than the two other groups at baseline (Control – Glucose difference ± SE: -0.11 ± 0.04; t = -2.77, P = 0.017; Glucose - Methylglyoxal: 0.112 ± 0.038; t = 2.907, p = 0.011).

Within sexes, nearly the same effect was observed for both sexes, with higher d-ROM levels at the end of the experiment than at Baseline and midpoint, and females also exhibited higher levels in February than at baseline (See **Table ESM2.2** and **Figure ESM2.1**). Furthermore, in February, females had higher d-ROM levels than males (difference ± SE: 0.148 ± 0.042; t = 3.542, P = 0.0005). These patterns were equally reflected by positive significant effects of longitudinal age on all groups and sexes (effect ± SE: 0.41 ± 0.109; t = 3.747, P = 0.0003).

When considering the interaction between treatment and month, a significant positive relationship between OXY and d-ROMs was found only for glucose at both baseline (slope ± SE: 0.003 ± 0.001; t = 0.001, P = 0.005) and in February (0.005 ± 0.001; t = 0.002, P = 0.007), for control birds in August (0.001 ± 0.0004; t = 0.0004, P = 0.002) and for methylglyoxal-supplemented birds at baseline (0.002326 ± 0.000538; t = 0.001, P = 0.003). The levels of d-ROMs were not significantly repeatable (R = 0.082, SE = 0.086, CI_95_[0, 0.287], P = 0.215). All results are shown in UCARR units.

#### Protein carbonyl (Oxidative Damage to Proteins)

Plasma protein carbonyl levels (nmoles/mg of protein) were significantly lower at the end of the experiment for all groups compared to their respective levels at all other months (**Figure 1.C.** and T**able ESM2.3**). This was reflected by a negative correlation with the longitudinal age component (slope ± SE: -0.246 ± 0.098, t = -2.51, P = 0.014). Moreover, protein carbonyl levels were higher in males (estimate ± SE: 0.055 ± 0.027; t = 2.007, P = 0.048). A decrease in protein carbonyl levels with increasing OXY was observed (slope ± SE: -0.001 ± 0.0002, t = -2.832, P = 0.005; **Figure ESM2.2**). Repeatability was moderate but significant (R = 0.213, SE = 0.113, CI_95_[0, 0.436], P = 0.014).

#### DNA damage

DNA damage, assessed as the percentage of 8-OH d-Guanosine relative to total DNA (**Figure 2.A**), showed a significant increase with longitudinal age in methylglyoxal-treated birds (slope ± SE: 0.507 ± 0.209, CI_95_[0.092, 0.921]). Birds in the methylglyoxal group moreover exhibited significantly higher levels at the end of the experiment than both control and glucose-supplemented birds (Control - Methylglyoxal difference ± SE: -0.98 ± 0.172, t = -5.706; P < 0.0001; Glucose - Methylglyoxal: -0.52 ± 0.205; t = -2.533, P = 0.032). They also showed higher levels than its own earlier measurements (Baseline - August: -0.461 ± 0.151; t = -3.044; P = 0.008; February - August: -0.531 ± 0.155; t = -3.428, P = 0.002). The control group showed lower levels at the end of the experiment than in February (February - August: 0.535 ± 0.171; t = 3.124, P = 0.006) and baseline (Baseline - August 0.45 ± 0.157; t = 2.86, P = 0.013). A significant increase with longitudinal age was also found in females (slope ± SE: 0.431 ± 0.216, CI_95_[0.004, 0.857]). There was no repeatability for this trait (R = 0, SE = 0.051, CI_95_[0, 0.173], P = 0.5).

**Figure 2.**
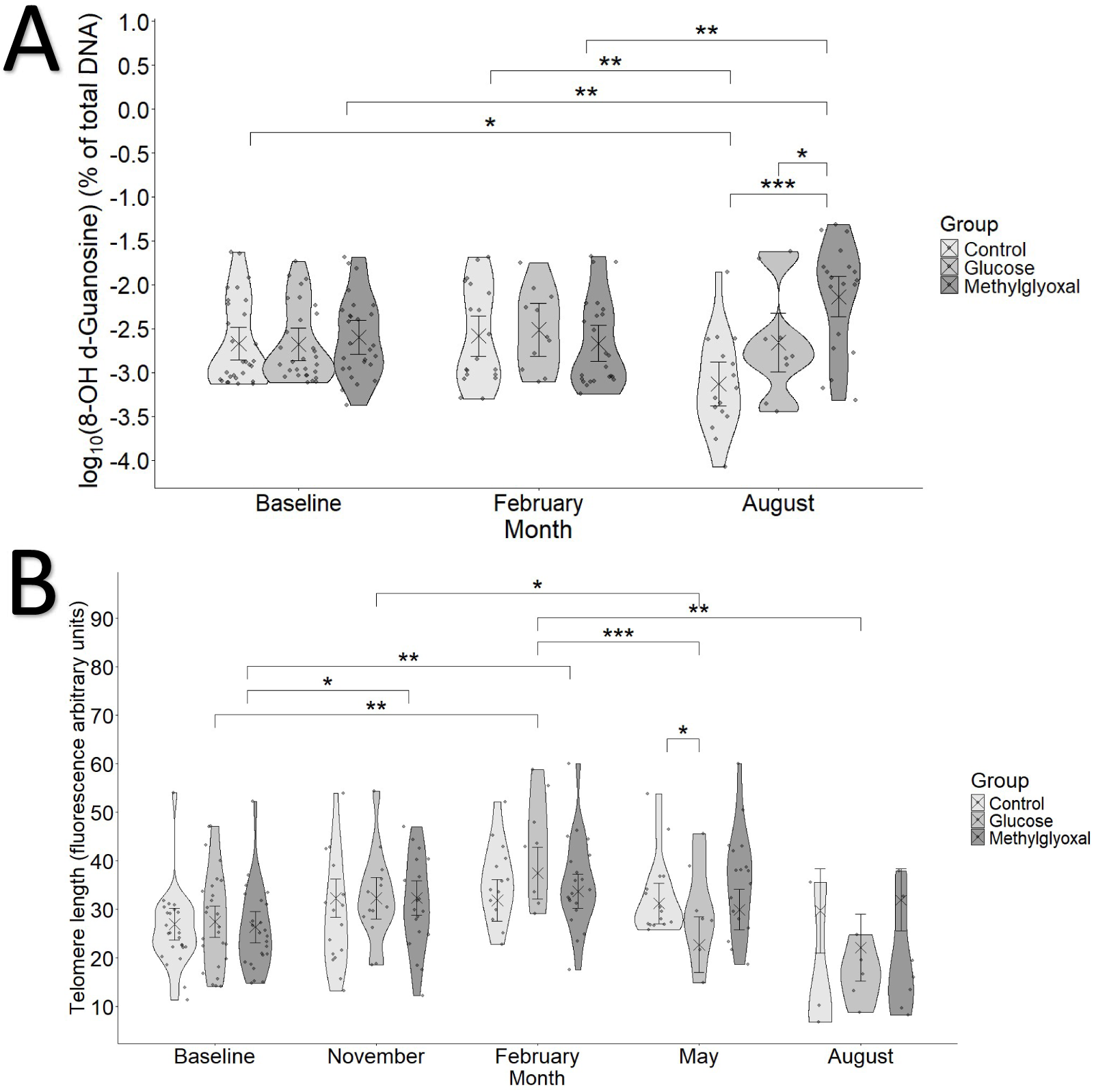
DNA damage (**A**) measured as 8 OH d-Guanosine and Telomere length (**B**) along the experiment and across treatment groups. Crosses and error bars represent model-estimated marginal means ±95% CI. Significance annotations are based on pairwise contrasts performed separately within months and within treatments.

#### Telomere Length and dynamics

As a general pattern, telomere length was significantly lower in older birds (horizontal age component; slope ± SE: -2.5015 ± 0.6307; t = -3.966, P = 0.0002). Telomere length also positively correlated with total sample DNA amount (log_10_ transformed, slope ± SE: 26.509 ± 3.671; t = 7.222, P = 9.7*10^-12^), meaning that telomere length was proportionally higher in samples with a low DNA quantity.

Seasonal increase in telomere length followed by a decrease can be inferred, although with differences between treatments (**Figure 2.B.**). Both supplemented groups showed a significant difference (i.e. longer telomeres) in February compared to baseline (Glucose: Baseline - February difference ± SE: -10.013 ± 2.86; t = -3.502, P = 0.005; Methylglyoxal: Baseline - February: -7.361 ± 2.1; t = -3.502, P = 0.005), with the methylglyoxal-treated birds already showing longer telomeres in February than at baseline (Baseline - November: -5.982 ± 2.05; t = -2.913, P = 0.033). In the glucose group, this increase was followed by a significant decrease in May (February - May: 14.781 ± 3.55; t = 4.159, P = 0.0005; November - May: 9.594 ± 3.36; t = 2.853, P = 0.038), resulting in values that were lower than those of control birds at that time (Glucose - Control: -8.505 ± 3.5; t = 2.431, P = 0.042), and a further decline was seen in August (February - August 15.358 ± 4.19; t = 3.663, P = 0.003), suggesting stronger seasonality in telomere length dynamics in the glucose group.

A significant positive effect of DNA damage on telomere length was found only for males (estimate ± SE: 351.8 ± 123; CI_95_[106, 598]), but there was no effect of DNA damage (assessed on the previous measurement session) on the telomere dynamics. Older individuals (by age at the beginning of the experiment; estimate ± SE: -0.273 ± 0.127; t = -2.152, P = 0.037) and longer telomeres before the change (-0.833± 0.046; t = -17.948, P <2*10^-16^) led to lower increase or higher decrease. On average, the change from baseline to February was positive, indicating telomere length gain (marginal mean ± SE: 0.696 ± 0.175; CI_95_[0.344, 1.049]), whereas the change from February to August was negative, indicating telomere length loss (-1.393 ± 0.292; CI_95_[-1.981, -0.806]). Telomere length repeatability was considerable and very significant (R = 0.362, SE = 0.084, CI_95_[0.19, 0.503], P = 6.14*10^-8^).

#### Cell Apoptosis

Apoptosis probability was significantly higher for all groups in November than in all later months, and the same was true in February, both at time 0 and after 24 hours, while the control group showed an increase in apoptosis probability from May to August (**Table ESM2.3**). Apart from November, when methylglyoxal-treated birds showed significantly lower apoptosis levels than both control and glucose-supplemented birds (**Figure 3**; Control / Methylglyoxal odds ratio ± SE: 2.248 ± 0.304; Z = 6, P < 0.0001; Glucose / Methylglyoxal: 1.769 ± 0.289; Z = 3.493, P = 0.001), treated birds tended to have higher apoptosis probability later in the experiment: both supplemented groups showing higher apoptosis probability than the control group in February (Control / Glucose: 0.321 ± 0.055; Z = -6.659, P < 0.0001; Control / Methylglyoxal: 0.46 ± 0.063; Z = -5.652, P < 0.0001) and May (Control / Glucose: 0.626 ± 0.122; Z = -2.409, P = 0.042; Control / Methylglyoxal: 0.266 ± 0.04; Z = -8.85, P < 0.0001), and methylglyoxal-supplemented birds showing higher levels than the glucose-supplemented birds in May (Glucose / Methylglyoxal 0.425 ± 0.08; Z = -4.571, P < 0.0001). Apoptosis levels in the methylglyoxal group were also higher than in both other groups in August (Control / Methylglyoxal: 0.409 ± 0.063; Z = -5.785, P < 0.0001; Glucose / Methylglyoxal: 0.565 ± 0.107; Z = -3.016, P = 0.007).

**Figure 3.**
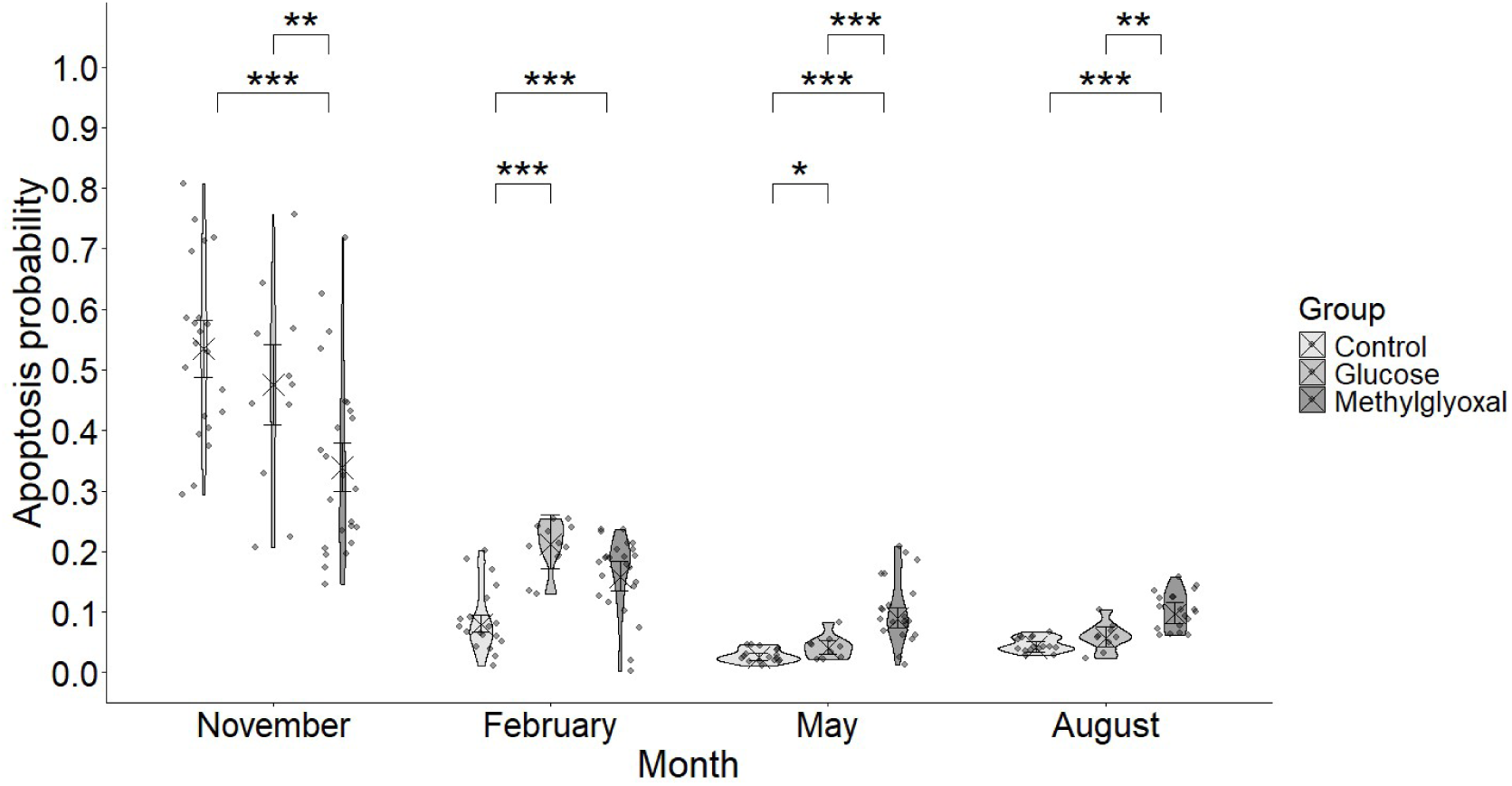
Apoptosis levels represented as a proportion of early apoptotic versus living cells along the experiment and across the treatment groups at the first measure (T0). The second measure, after 24 hours (T24), is shown in **ESM2**. Crosses and error bars represent model-estimated probabilities ±95% CI. Significance annotations are based on pairwise contrasts performed separately within months. Comparisons within treatments across months are not included here to improve the figure visibility, due to their high number of significant results, with November and February having higher levels than all later month for all groups.

After 24 hours, apoptosis levels were significantly higher (effect ± SE: 0.166 ± 0.006; Z = 26.28, P < 2*10^-16^), but apoptosis appeared only affected by methylglyoxal treatment (based on within month contrasts) with higher levels than in both other groups in May (**Figure ESM2.3,** Control / Methylglyoxal odds ratio ± SE: 0.633 ± 0.094; Z = -3.062, P = 0.006; Glucose / Methylglyoxal: 0.55 ± 0.103; Z = -3.194, P = 0.004) and August (Control / Methylglyoxal : 0.652 ± 0.101; Z = -2.769, P = 0.015; Glucose / Methylglyoxal: 0.633 ± 0.12; Z = -2.415, P = 0.042), and lower levels in November (Control / Methylglyoxal: 2.205 ± 0.298; Z = 5.856, P < 0.0001; Glucose / Methylglyoxal: 1.758 ± 0.287; Z = 3.456, P = 0.002).

These patterns are confirmed by a significant decrease in apoptosis probability with longitudinal age (slope ± SE: -5.159 ± 0.362, z = -14.255, P < 2*10^-16^), which was lower in the methylglyoxal group at T0 and T24 (**Table ESM2.4**). Given the significant positive effect of the interaction between longitudinal and horizontal age components (estimate ± SE: 0.15 ± 0.074, z = 2.015, P = 0.044), apoptosis levels decreased less over the experiment in older birds. However, the horizontal age component had a positive significant effect only when not accounting for longitudinal age (effect ± SE: 0.063 ± 0.031; Z = 2.03, P = 0.043), indicating older birds would have higher apoptosis.

Finally, although the interaction of sex with time (i.e. higher effects for males after 24 hours: 0.031 ± 0.009; Z = 3.42, P = 0.001) and some other interactions (i.e. Time:Month:Sex, Time:Treatment:Sex and Time:Month:Treatment:Sex) were found significant, no significant differences between sexes were found for any of the possible subgroups after performing marginal means contrasts. Finally, apoptosis probability increased after 24 hours for all groups, except for a decrease in the glucose-supplemented birds in February and the methylglyoxal group in August, and a non-significant change for the glucose group in August (**Table ESM2.5**).

#### Mitochondrial superoxide production

Mitochondrial superoxide production patterns were different between living and early-apoptotic cells, with a higher intercept in the early apoptotic main model (de-transformed intercept ± SE: 19,098.53 ± 1.107 MitoSox fluorescence units) than in the living cells main model (de-transformed intercept ± SE: 2,312.065 ± 1.12 MitoSox fluorescence units).

##### Living cells

Living cells showed significantly higher superoxide production levels in November than in later months, with the lowest values in February, followed by an increase in May and a slight decrease in August. However, this last decrease is not significant in the 24-hour measurement (**Table ESM2.6**). Within months, the only significant difference was found in May (**Figure 3.A**), when the methylglyoxal-supplemented group had a significantly lower superoxide production than the control (Control - Methylglyoxal difference ± SE: 0.664 ± 0.079; t = 8.386, P < 0.0001) and glucose-treated group (Glucose - Methylglyoxal: 0.667 ± 0.094; t = 7.105 P < 0.0001), with the same results after 24 hours (**Figure ESM2.4.A;** Control - Methylglyoxal: 0.664 ± 0.079; t = 8.386, P < 0.0001; Glucose - Methylglyoxal: 0.667 ± 0.094; t = 7.105, P < 0.0001).

A significant relationship between superoxide production and the number of living cells was found only in February (marginal slope ± SE: -8.23*10^-5^ ± 7.42*10^-6^; CI_95_[-9.69*10^-5^, -6.77*10^-5^]) and May (-6.29*10^-5^ ± 1.34*10^-5^; CI_95_[-8.93*10^-5^, -3.65*10^-5^]) for initial (T0) measures, and in November (8.74*10^-5^ ± 8.70*10^-6^; CI_95_[7.02*10^-5^, 1.04*10^-4^) for 24-hour (T24) measures, although with opposite directions of effect (negative at T0 and positive at T24). Finally, a significant positive longitudinal age effect was observed (effect ± SE: 126.5 ± 16.25; t = 7.782, P = 3.22*10^-13^), and superoxide levels were significantly higher after 24 hours only in February (T0 - T24 difference ± SE: -0.335 ± 0.039; t = -8.479, P < 0.0001) and August (-0.287 ± 0.042; t = - 6.796, P < 0.0001). Repeatability for superoxide production in living cells was relatively low but very significant (R = 0.118, SE = 0.054; CI_95_[0.006, 0.227], P = 1.59*10^-12^).

##### Early apoptotic cells

In the case of early apoptotic cells, the main pattern observed was a lower superoxide production in the treated birds, i.e. in the glucose (difference ± SE: Control - Glucose: 0.265 ± 0.066; t = 4.027, P = 0.0002) and methylglyoxal (Control - Methylglyoxal: 0.318 ± 0.05; t = 6.315, P < 0.0001) groups compared to the control group in February. All groups showed a reduction compared to their previous levels in November (difference ± SE: Control: November - February: 0.286 ± 0.078; t = 3.675, P = 0.002; Glucose: November - February: 0.597 ± 0.081; t = 7.399, P < 0.0001; Methylglyoxal: November - February: 0.687 ± 0.057; t = 12.124, P < 0.0001), and this increased again in May (Control: February - May: -0.475 ± 0.142; t = -3.339, P = 0.005; Glucose: February - May -0.76 ± 0.146; t = -5.19, P < 0.0001; Methylglyoxal: February - May: -0.659 ± 0.105; t = -6.303, P < 0.0001; **Figure 4.B**). These effects were similar after 24 hours (**Figure ESM2.4.B**; within month effects: difference ± SE: Control - Glucose: 0.265 ± 0.066; t = 4.027, P = 0.0002; Control - Methylglyoxal: 0.318 ± 0.05; t = 6.315, P < 0.0001), except that only the glucose and methylglyoxal groups showed a decrease in February (Glucose: November - February: 0.305 ± 0.083; t = 3.675, P = 0.002; Methylglyoxal: 0.394 ± 0.057; t = 6.945, P < 0.0001) followed by a later increase in May (Glucose February - May: -0.573 ± 0.129; t = -4.447, P = 0.0001; Methylglyoxal: -0.472 ± 0.086; t = -5.467, P < 0.0001).

**Figure 4.**
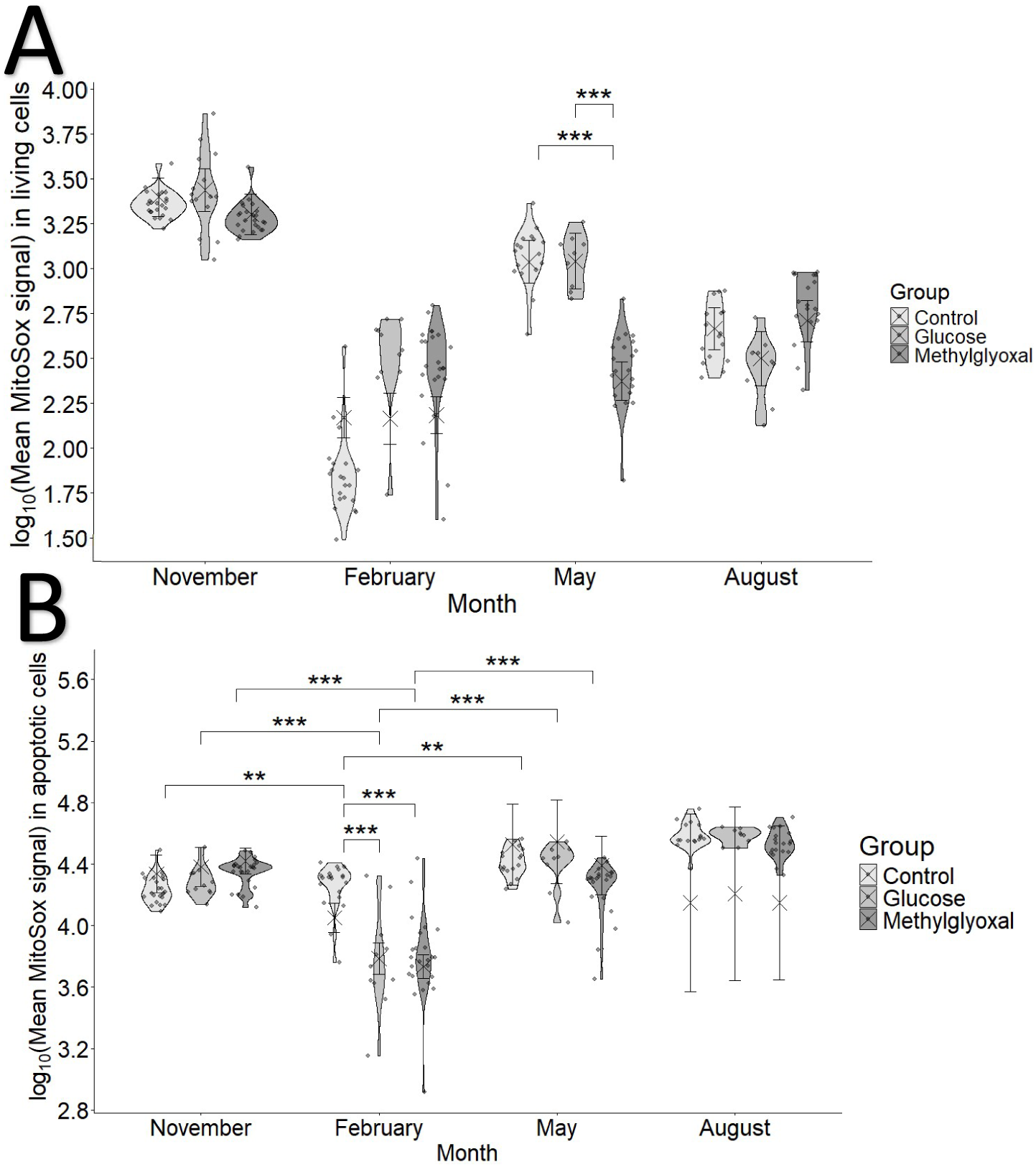
Mitochondrial superoxide production of (**A**) living cells and (**B**) apoptotic cells, represented as the decimal logarithm of MitoSox fluorescence signal along the experiment and across the treatment groups at the first measure (T0). The second measure, after 24 hours (T24), is shown in **ESM2**. Crosses and error bars represent model-estimated marginal means ±95% CI. Significance annotations are based on pairwise contrasts performed separately within months and within treatments. As in Figure 2, within group (across months) comparisons are excluded from **A** in order to improve visibility.

For early apoptotic cells, a significant negative relationship between the number of cells and superoxide production was found in November (slope ± SE: -1.79*10^-5^ ± 7.06*10^-6^, CI_95_[-3.18*10^-5^, -3.96*10^-6^]) and February (-9.54*10^-5^ ± 1.89*10^-5^, CI_95_[-1.33*10^-4^, -5.82*10^-5^]). No significant effects of longitudinal age were found, neither for its linear nor quadratic component. Finally, after 24 hours, the superoxide production was significantly lower in early apoptotic cells in November (T0 - T24 difference ± SE: 0.063 ± 0.03, t = 2.067, P = 0.04) and higher in February (: - 0.229 ± 0.032; t = -7.207, P < 0.0001) and August (-0.074 ± 0.035; t = -2.117, P = 0.035). Repeatability was very low but highly significant for superoxide production in early apoptotic cells (R = 0.031, SE = 0.038, CI_95_[0, 0.128], P = 6.93*10^-6^).

##### Superoxide temporal dynamics

Regarding superoxide production dynamics, (i.e. the difference between T0 and T24), the control group showed a decrease in every month except August, in both living (November marginal mean ± SE: -1.417 ± 0.192, CI_95_[-1.7944, -1.039]; February: -0.606 ± 0.201, CI_95_[-1.001, -0.21]; May: -0.604 ± 0.223, CI_95_[-1.043, -0.164]) and early apoptotic cells (November: -0.587 ± 0.292, CI_95_[-1.163, -0.011]; February: -1.46 ± 0.306, CI_95_[-2.062, -0.857]; May: -0.741 ± 0.341, CI_95_[-1.412, -0.07]). In contrast, glucose-supplemented birds increased superoxide production in February for both living (2.597 ± 0.277, CI_95_[2.05, 3.143]) and apoptotic cells (2.266 ± 0.423, CI_95_[1.433, 3.1]), but decreased in November only for living cells (-1.16 ± 0.237, CI_95_[-1.627, - 0.692]). Methylglyoxal-treated individuals also showed an increase for both living (2.248 ± 0.188, CI_95_[1.878, 2.618]) and apoptotic cells (1.8875 ± 0.286, CI_95_[1.324, 2.451]) in February and in May (living: 1.727 ± 0.201, CI_95_[1.332, 2.122]; apoptotic: 0.816 ± 0.306, CI_95_[0.213, 1.42]), while decreasing in November (living: -1.966 ± 0.188, CI_95_[-2.336, -1.597]; early apoptotic: -0.825 ± 0.286, CI_95_[-1.389, -0.261]). Finally, the difference increased with longitudinal age in the control (slope ± SE: 2.202 ± 0.667, CI_95_[0.884, 3.52]) and methylglyoxal groups (1.883 ± 0.61, CI_95_[0.678, 3.09]), only for living cells.

#### Correlation with additional physiological parameters and effects on mortality hazard

Models testing the effects of other variables measured in these birds, as reported in Moreno Borrallo et al. 2026 (i.e. metabolic rate, circulating glucose and glucose-derived damage markers), on oxidative status and vice versa (see **ESM2 Box 2**), yielded mixed results. First, models testing the effects of oxygen consumption rates (VO_2_ in mL/min) on mitochondrial superoxide (O ^·-^) production (either for living or early-apoptotic cells), and on oxidative stress (d-ROMs, protein carbonyls or DNA damage), did not shown any significant effects.

Whole blood glucose levels showed a positive correlation with d-ROMs only in males (marginal slope ± SE: 0.658 ± 0.229; CI_95_[0.207, 1.109]), whereas plasma glucose showed a negative correlation with d-ROMs in females during February (marginal slope ± SE: -2.716 ± 0.699; CI_95_[-4.098, -1.33]) and a positive effect in protein carbonyl in males (1.226 ± 0.53; CI_95_[0.175, 2.278). No significant effects of oxidative stress on albumin glycation levels were found in any of the three performed models (d-ROMs, protein carbonyls and DNA damage on glycation), although a positive relationship between OXY and glycation levels were found in the models testing for protein carbonyl (slope ± SE: 0.009 ± 0.004; t = 2.225, P = 0.029) and DNA damage effects (0.009 ± 0.004; t = 2.385, P = 0.019). Regarding reactive dicarbonyl compounds, only a positive significant association of glyoxal with d-ROMs levels in males was found (marginal slope ± SE: 0.264 ± 0.118, CI_95_[0.029, 0.4985]). Regarding AGE, plasma CML levels had a positive significant correlation with OXY in the model testing for effects of d-ROMs (slope ± SE: 4.843*10^-5^ ± 2.261*10^-5^; t = 2.142, P = 0.035), while CEL levels were correlated with DNA damage, with a significant negative relationship in males (marginal slope ± SE: -0.022 ± 0.01; CI_95_[-0.043, - 0.001]). Finally, a negative significant effect of protein carbonyl levels (coefficient ± SE: β= -0.31 ± 0.12, Z = -2.575, P = 0.01) and its interaction with age on mortality hazard was found (coefficient ± SE: β= -0.179 ± 0.074, Z = -2.406, P = 0.016). No effect of any of the other oxidative status variables or telomere length was found on mortality hazard.

## Discussion

### Methylglyoxal supplementation increases DNA damage without impacting survival

The increase in DNA damage observed exclusively in the methylglyoxal-treated group, and not in the glucose group, may be attributed to the low intracellular exposure of avian erythrocytes to glucose (Johnstone et al. 1998). However, the lack of significant repeatability for DNA damage suggests cautious interpretation, particularly when compared to protein carbonyls, which showed moderate repeatability. The repeatability patterns for OXY and d-ROMs align with findings in wild collared flycatcher (*Ficedula albicollis*, Récapet et al. 2019), but contrast with previous zebra finch studies (Beamonte-Barrientos and Verhulst 2013). A recent study in chicken showed that intraperitoneal methylglyoxal injection induces oxidative damage to lipids in liver and pectoralis muscle, but not in plasma (Okino et al. 2026), which may explain why we only found damage in cells. Instead, they show an increase in plasma antioxidant capacity, contrary to what we see here, which may represent a difference between an acute and a chronic response.

None of the oxidative status parameters affected survival, except for a negative effect of protein carbonyl levels on mortality hazard, both alone and in interaction with age. This suggests that the apparently protective effect of plasma protein carbonyl levels on mortality hazard increases with age. This finding is difficult to explain, as we controlled for protein concentration, ruling out metabolic activity as a confounding factor (e.g., higher protein anabolism levels). Our results contrast with a previous study showing increased protein carbonyl and 8-OH-d-Guanosine levels in captive zebra finches after one year, even in controls, with an impact on survival of DNA damage (Marasco et al. 2017). The differences in the way they measure damage, i.e. protein carbonyl in cells vs. DNA damage in plasma (the opposite of ours) may explain these discrepancies, as plasma 8-OH-d-Guanosine could reflect cell death and DNA release, a worse scenario than DNA damage found in cells, while cellular protein damage may be more deleterious, given the limited protein synthesis in bird erythrocytes (Sinclair and Brasch 1975). Finally, they showed that the effects on age-related damage accumulation and of this damage on survival is context-dependent, with harsh foraging environment suppressing survival effects (see also Constantini 2014 for a review on oxidative stress effects on the wild).

### Oxidative damage and antioxidant relationships: response or protection?

Interestingly, despite the lack of effects of glucose supplementation on oxidative status, the glucose-supplemented group exhibited elevated d-ROMs levels at baseline, perhaps linked to higher baseline whole blood (but not plasma) glucose levels (Moreno Borrallo et al. 2026). Previous studies in birds, such as house sparrows (*Passer domesticus*), have shown that blood glucose levels positively relate to oxidative damage markers like malondialdehyde (Vágási et al. 2020), although this pattern is not consistent across songbird species (Vágási et al. 2024).

Oxidative damage levels may be influenced by both reactive oxygen species (ROS) production (see below) and antioxidant defences. In plasma, OXY showed a significant positive relationship with d-ROMs only in the supplemented groups at baseline, the glucose group in February, and the control group in August. This pattern could reflect a defensive response where antioxidant levels rise in response to increased oxidative damage (Costantini and Verhulst 2009). This scenario is partly supported by the mentioned effects being found in the glucose-supplementation group precisely when they showed higher d-ROMs levels (i.e. baseline and February), or in the control group in August, a period where oxidative damage was also generally increased. It is also in line with the positive relationship of OXY with glycated albumin levels and plasma CML, all compounds that reflect oxidative stress-induced damage. However, this interpretation is complicated by the lack of a consistent relationship across all groups and time points. For instance, the methylglyoxal group showed this relationship at baseline without a corresponding increase in damage levels.

Conversely, plasma antioxidant levels exhibited a significant negative relationship with protein carbonyl levels (regardless of treatment), suggesting that a lower antioxidant protection may lead to higher oxidative damage. The relationship between oxidative damage and glycation was model-dependent, highlighting the complexity of these interactions. Such variability in the relationships between oxidative damage and defence is frequently reported in the literature and may arise from variation in the nutritional or energetic states that modulate responses (Costantini and Verhulst 2009; Costantini 2019). For example, the season-dependent negative effect of plasma glucose on d-ROMs levels in females (where higher plasma glucose predicted lower organic hydroperoxide levels) suggests that environmental conditions (see weather variation on **ESM3**), or an associated variation in food intake (Meijer et al. 1996; Geiger et al. 2012) and metabolism (Bauchinger et al. 2010; Campbell, et al. 2025) may interact with oxidative status. However, we did not observe an effect of metabolic rate, which increased in winter in the supplemented birds (Moreno Borrallo et al. 2026), on oxidative status. Sex dependent effects also complicate the issue, as a significant effect of whole blood glucose on d-ROMs was only observed in males. Additionally, plasma glucose levels positively correlated with plasma protein carbonyl content, a marker related to glycation (reviewed in Mitsugu 2020), only in males, independently of the treatment.

### Metabolism does not influence oxidative damage

Whole-body oxygen consumption did not affect oxidative status in our study. This opposes previous findings in zebra finches, where increase in metabolic rate induced by acute cold exposure did not have an effect on OXY and d-ROMs levels in zebra finches (Beamonte-Barrientos and Verhulst 2013). Similarly, in another study by Stier et al. (2014), neither the increase in metabolic rate induced by acute nor by chronic cold-exposure showed an increase in oxidative damage in plasma, although they showed a decrease in OXY with chronic cold exposure and an increase in DNA damage with acute, but not with chronic cold exposure, which would be more comparable to our case, as our birds were exposed to outdoor low temperatures during winter (see **ESM3**).

The complex and diverse results may reflect tissue-specific effects (Costantini 2019), or the need to account for factors such as the mitochondrial uncoupling state (Hou et al. 2021; Stier et al. 2014). More integrative metabolic measures, such as Standard Metabolic Rate (SMR) or Daily Energy Expenditure (DEE), might better capture ecologically relevant constraints. For instance, summit metabolic rate, but not basal metabolic rate, has been linked to winter survival in black-capped chickadees (*Poecile atricapillus*, Petit et al. 2017). Future studies should consider these measures to improve ecological realism (Briga and Verhulst 2017).

### Seasonal variation of telomere length is enhanced by glucose supplementation: a potentially antagonistic mechanism?

The negative effect of the horizontal age component on telomere length suggests that long term measurements are necessary to detect telomere attrition, as no clear longitudinal shortening pattern emerged within a single year. Instead, we observed a seasonal cycle of telomere lengthening during winter and shortening during spring. Telomere dynamics models confirmed an increase from baseline to February and a decrease from February to August. Interestingly, seasonal telomere length variation has previously been reported in mammals (Criscuolo et al. 2020; Redon et al. 2024), although sometimes possibly due to cell type variation (Beaulieu et al. 2017; Rehkopf et al. 2014). Our results are, to our knowledge, the first time a seasonal pattern is reported in birds.

Glucose-supplemented birds presented more pronounced seasonal telomere dynamics, with greater winter lengthening followed by a faster spring shortening. This suggests that glucose supplementation may have antagonistic effects on telomeres, offering short-term advantages but with a potential long-term cost, especially given that telomere dynamics has been related with lifespan in zebra finches (i.e. higher telomere attrition predicts lower lifespan; Tangili et al. 2026). This seasonal pattern may also relate to the increased resting metabolic rate during winter in treated birds, which persisted longer in the glucose-supplemented group (Moreno Borrallo et al. 2026). However, as the supplementation was maintained over time during the whole duration of our experiment, we cannot distinguish long-term carryover effects from chronic exposure effects. For this, future studies should explore the impact on telomeres of shorter supplementation periods at different times, including the effects during or shortly after the supplementation and those that appear later on, to clarify these dynamics.

Finally, DNA damage positively correlated with telomere length measured simultaneously to it only in males. This adds to the abovementioned male-restricted oxidative damage effects, although again the support is only correlational. Nevertheless, damage did not affect telomere dynamics assessed in the period immediately after such damage measurement. Since increases in DNA damage by a treatment were only observed at the end of the experiment, we could not assess whether the supplementation impacted telomere dynamics through DNA damage beyond mere correlation. A recent meta-analysis demonstrating the effects of oxidative stress on telomeres in vivo (Armstrong and Boonekamp 2023) highlighted that the technique used for telomere length assessment influences observed effects, with Telomere Restriction Fragments (TRF) being more likely to point out a significant correlation than qPCR. Unfortunately, qFISH measures were not included in the meta-analysis, likely due to a lack of studies using this technique in the context of oxidative status effects on telomeres *in vivo*. Seasonal changes in telomerase activity should also be considered in the future as a potential underlying mechanism mediating the observed effects. However, telomerase expression in bone marrow and blood is apparently low in adult zebra finches (Haussmann et al. 2004).

### Apoptosis is induced by glucose and more markedly by methylglyoxal supplementation

Apoptosis levels were highest and very variable in November, with methylglyoxal-supplemented birds initially showing lower levels than control and glucose groups. This variability may reflect measurement errors or higher overall cell death during this high mortality period (Moreno Borrallo et al. 2026). Later, the pattern was reversed, with methylglyoxal-supplemented birds exhibiting higher proportions of apoptotic red blood cells than controls in all subsequent months, and higher levels than glucose-supplemented birds, except in February. Glucose supplementation also increased apoptosis probability in February, at a similar level than methylglyoxal, and in May, though not in August. These results are generally in line with expectations, particularly for methylglyoxal, which exhibited more deleterious effects on cell survival, consistent with findings in human cells (review in Vašková et al. 2025). The more pronounced effects of methylglyoxal compared to glucose may relate to the low permeability of avian red blood cells to glucose (Johnstone et al. 1998), enabling a greater resistance to glucose-associated damage. The increase in apoptosis with horizontal age suggests an age decline in cell resistance, similar to patterns observed for oxidative stress resistance in other bird species (Devevey et al. 2010; Bize et al. 2014). In zebra finches, an age-related decrease in haematocrit has been reported, though it seems unrelated to erythrocyte number (Coughlan et al.2022), leaving unclear whether our age-related increase in apoptosis contributes to this pattern.

The lack of increase in apoptosis levels 24 hours after the initial measurement, or even the reversed pattern, in treated birds towards the end of the experiment (August 2023), may indicate higher resilience after prolonged exposure or selective disappearance of the lower-quality individuals, particularly in the glucose group, which showed no difference from the controls in August 2023.

### Mitochondrial superoxide production and apoptosis: partial evidence for a direct link in supplemented birds

#### Seasonal Effects and Treatment-Specific Patterns

We observed a transient decrease in superoxide production during winter for both living and apoptotic cells, particularly in supplemented birds for the latter. This pattern may be attributed to increased mitochondrial uncoupling due to elevated thermogenesis, as resting metabolic rate (RMR) was higher in treated birds during this period of cold ambient temperatures (Moreno Borrallo et al., 2026), although it is unclear if such process would be observed in red blood cells, as the increase in metabolism was assessed in whole body. While methylglyoxal and glucose supplementation may facilitate mitochondrial uncoupling —a mechanism known to mitigate cold-induced oxidative stress (Stier et al., 2014a & b)— this does not directly translate to decreased apoptosis. Instead, the transient decrease in superoxide production during winter suggests a potential plastic adaptive response to cold stress. Alternatively, a more direct impact of the treatments on mitochondrial functioning could be triggered by methylglyoxal, and needs to be further explored (de Souza Prestes et al. 2022).

#### Dynamics of Superoxide Production and Apoptosis

When examining the ex vivo dynamics of superoxide production (differences between initial and 24-hour measurements), we found that superoxide levels increased in treated birds for both apoptotic and living cells in February, and later in May for methylglyoxal-supplemented birds. This pattern, combined with a decrease in superoxide production after 24 hours for the methylglyoxal group in November, aligns partly with results in apoptosis levels, suggesting that ex vivo superoxide dynamics is a better predictor of apoptosis than raw values. Nevertheless, the apoptosis levels after 24 hours were not higher than the control in the treated birds in February, which, indicates that alternative pathways may also play a role in explaining the observed apoptosis patterns.

#### Limitations and Future Directions

A key limitation of our study is the low individual repeatability of the MitoSox signal (∼12% for living cells and ∼0.3% for early apoptotic cells). MitoSox, while useful for assessing mitochondrial oxidant status, may not exclusively represent mitochondrial superoxide production (Zielonka and Kalyanaraman, 2010). Additionally, we did not assess mitochondrial coupling status, which complicates the interpretation of the relationship between oxygen consumption and superoxide production (Hou et al., 2021; Stier et al., 2014a, 2014b). Indeed, Dawson and Salmon (2020) showed a cross-sectional age-related decline in red blood cell leak state per unit of mitochondria in zebra finches, perhaps implying a higher ROS production. Nevertheless, we did not find any age-related change in mitochondrial superoxide production, as previously reported (also cross-sectionally) in zebra finches liver and skeletal muscle (Salmon et al., 2022).

## Conclusion

Mitochondrial superoxide production is a well-established contributor to cell death across various biological models (Stowe and Camara, 2009; Kanamori et al., 2010; Ma et al., 2011; Fujii et al., 2022). In our study, apoptotic cells consistently exhibited higher superoxide levels than living cells, confirming the role of superoxide in apoptosis. Moreover, our results suggest that ex vivo mitochondrial superoxide production dynamics, rather than its raw levels, may explain the increased apoptosis observed in methylglyoxal- and glucose-supplemented birds.

However, the transient seasonal effects and the lack of a full consistency in the relationship between superoxide production and apoptosis in treated birds indicate that other mechanisms—such as direct glycoxidative damage or metabolic shifts—may also play a role in driving apoptosis. Further research is needed to elucidate these mechanisms and their interactions with mitochondrial function.

## Supporting information

ESM1 - Birds list

ESM2 - Extra results

ESM3 - Weather

## Acknowledgements

This research was funded by the Agence Nationale de la Recherche (ANR, project AGEs, ANR21-CE02-0009), and the CNRS/University of Toronto Joint Call 2021. RCC has received funding from both the European Union’s Horizon 2020 research and innovation programme under the Marie Skłodowska-Curie grant agreement No. 101026888 and the Valencian Government under the CDEIGENT grant [2023/29]. We thank Helène Gachot-Neveu, David Bock, Aurélie Hranitzky and Nicolas Spanier for their work in the animal facility, Yaël Metzger for help on part of the baseline data collection and Sandrine Zhan for the help in DNA extraction. We also thank Max Planck Institute for Biological Intelligence (Pöcking-Seewiesen, Bavaria, Germany) and Bielefeld University (Bielefeld, Germany) for providing part of the birds used in this study.

## Ethics statement

This study adhered all the legal and ethical regulations and was authorized by the French Ministry of Secondary Education and Research, APAFIS #32475-2021071910541808 v5.

