## Supplementary material for "Contrasting effects of glucose and methylglyoxal supplementation on blood oxidative status, blood cells’ telomere dynamics and apoptosis in birds": ESM1 - Birds list

### ESM 1. Birds

#### Birds from the zebra finches' experiment

#### Provenance of the birds

**Table ESM1.1.** Zebra finches forming part of the experiment, with their origin.

| Bird_ID | Origin |
|---|---|
| 19 | Max-Planck-Institute for Ornithology |
| 35 | Max-Planck-Institute for Ornithology |
| 049 | Pet shop (2022) |
| 60 | Max-Planck-Institute for Ornithology |
| 086 | DEPE, originally from pet shop |
| 102 | Pet shop (2022) |
| 119 | DEPE, originally from pet shop |
| 127 | DEPE - Strasbourg |
| 132 | DEPE - Strasbourg |
| 139 | DEPE, originally from pet shop |
| 147 | DEPE, originally from pet shop |
| 154 | DEPE, originally from pet shop |
| 156 | DEPE, originally from pet shop |
| 165 | DEPE - Strasbourg |
| 177 | Pet shop (2022) |
| 618 | Bielefeld University |
| 840 | Bielefeld University |
| 1008 | Bielefeld University |
| 1052 | Bielefeld University |
| 010 K18 | Max-Planck-Institute for Ornithology |
| 013 K18 | Max-Planck-Institute for Ornithology |
| 019 K18 | Max-Planck-Institute for Ornithology |
| 020 K18 | Max-Planck-Institute for Ornithology |
| 022 K18 | Max-Planck-Institute for Ornithology |
| 029 K18 | Max-Planck-Institute for Ornithology |

| | |
|---|---|
| 035 K18 | Max-Planck-Institute for Ornithology |
| 044 K18 | Max-Planck-Institute for Ornithology |
| 045 K18 | Max-Planck-Institute for Ornithology |
| 047 K18 | Max-Planck-Institute for Ornithology |
| 049 K18 | Max-Planck-Institute for Ornithology |
| 053 K18 | Max-Planck-Institute for Ornithology |
| 054 K18 | Max-Planck-Institute for Ornithology |
| 059 K18 | Max-Planck-Institute for Ornithology |
| 061 K18 | Max-Planck-Institute for Ornithology |
| 071 K18 | Max-Planck-Institute for Ornithology |
| 073 K18 | Max-Planck-Institute for Ornithology |
| 076 K18 | Max-Planck-Institute for Ornithology |
| 079 K18 | Max-Planck-Institute for Ornithology |
| 088 K18 | Max-Planck-Institute for Ornithology |
| 089 K18 | Max-Planck-Institute for Ornithology |
| 099 K18 | Max-Planck-Institute for Ornithology |
| 10 16D | Max-Planck-Institute for Ornithology |
| 104 16D | Max-Planck-Institute for Ornithology |
| 104 S18 | Max-Planck-Institute for Ornithology |
| 106 16D | Max-Planck-Institute for Ornithology |
| 107 S18 | Max-Planck-Institute for Ornithology |
| 109 S18 | Max-Planck-Institute for Ornithology |
| 11 16D | Max-Planck-Institute for Ornithology |
| 112 K18 | Max-Planck-Institute for Ornithology |
| 115 S18 | Max-Planck-Institute for Ornithology |
| 119 16D | Max-Planck-Institute for Ornithology |
| 119 K19 | Max-Planck-Institute for Ornithology |
| 119 S18 | Max-Planck-Institute for Ornithology |
| 12 16D | Max-Planck-Institute for Ornithology |
| 121 K18 | Max-Planck-Institute for Ornithology |
| 127 K18 | Max-Planck-Institute for Ornithology |
| 129 K18 | Max-Planck-Institute for Ornithology |
| 132 S18 | Max-Planck-Institute for Ornithology |

| | |
|---|---|
| 135 K18 | Max-Planck-Institute for Ornithology |
| 135 K19 | Max-Planck-Institute for Ornithology |
| 137 K18 | Max-Planck-Institute for Ornithology |
| 145 K18 | Max-Planck-Institute for Ornithology |
| 145 S18 | Max-Planck-Institute for Ornithology |
| 148 S18 | Max-Planck-Institute for Ornithology |
| 151 K18 | Max-Planck-Institute for Ornithology |
| 153 K18 | Max-Planck-Institute for Ornithology |
| 158 K18 | Max-Planck-Institute for Ornithology |
| 173 16D | Max-Planck-Institute for Ornithology |
| 173 K19 | Max-Planck-Institute for Ornithology |
| 175 16D | Max-Planck-Institute for Ornithology |
| 194 16D | Max-Planck-Institute for Ornithology |
| 196 16D | Max-Planck-Institute for Ornithology |
| 205 16D | Max-Planck-Institute for Ornithology |
| 220 16D | Max-Planck-Institute for Ornithology |
| 222 16D | Max-Planck-Institute for Ornithology |
| 230 16D | Max-Planck-Institute for Ornithology |
| 234 16D | Max-Planck-Institute for Ornithology |
| 235 16D | Max-Planck-Institute for Ornithology |
| 242 16D | Max-Planck-Institute for Ornithology |
| 254 16D | Max-Planck-Institute for Ornithology |
| 27 16D | Max-Planck-Institute for Ornithology |
| 29 16D | Max-Planck-Institute for Ornithology |
| 295 K19 | Max-Planck-Institute for Ornithology |
| 37 16D | Max-Planck-Institute for Ornithology |
| 4 16D | Max-Planck-Institute for Ornithology |
| 70 16D | Max-Planck-Institute for Ornithology |
| 71 16D | Max-Planck-Institute for Ornithology |
| 78 16D | Max-Planck-Institute for Ornithology |
| 89 17S | Max-Planck-Institute for Ornithology |
| S520F/008 | Pet shop (2022) |

**Table ESM1.2.** Zebra finches forming part of the experiment, with their sex (assigned by visual inspection of the plumage), allocated treatment, age in years, body mass in grams and morphometric variables (in cm). All the measures presented here correspond to their values at the start of the experiment.

| Bird_ID | Sex | Treatment | Tarsus | Head_beak | Wing | Age_years | BM |
|---|---|---|---|---|---|---|---|
| 19 | female | Methylglyoxal | 14,5 | 24,3 | 57 | 5,884931507 | 14,60 |
| 35 | female | Methylglyoxal | 14,2 | 22,9 | 58 | 4,18630137 | 13,90 |
| 049 | female | Glucose | 15,3 | 26,1 | | | 18,20 |
| 60 | male | Control | 14,6 | 23,7 | 57 | 6,178082192 | 13,40 |
| 086 | male | Methylglyoxal | 14,4 | 24 | 59 | 4,016438356 | 16,30 |
| 102 | female | Control | 16,1 | 26,3 | | | 20,35 |
| 119 | female | Control | 17 | 26 | 62 | 4,726027397 | 21,90 |
| 127 | female | Control | 14,6 | 24,3 | 60 | 3,04109589 | 15,70 |
| 132 | male | Methylglyoxal | 15 | 25,7 | | 3,005479452 | 18,45 |
| 139 | male | Control | 14,65 | 23,95 | 60 | 2,969863014 | 15,30 |
| 147 | male | Glucose | 14,5 | 23,9 | 60 | 2,969863014 | 20,60 |
| 154 | male | Glucose | 14,9 | 24,1 | 58 | 2,969863014 | 15,70 |
| 156 | male | Glucose | 15,7 | 25,5 | 60 | 2,969863014 | 18,10 |
| 165 | male | Glucose | 14,8 | 23,3 | 58 | 2,649315068 | 12,20 |
| 177 | female | Glucose | 16,2 | 26,1 | 61 | | 19,50 |
| 618 | male | Control | 14 | 24,9 | 56 | 4,123287671 | 13,25 |
| 840 | male | Control | 14,5 | 23,1 | 53 | 3,761643836 | 11,35 |
| 1008 | male | Methylglyoxal | 13,9 | 23,5 | 56 | 2,183561644 | 11,95 |
| 1052 | male | Methylglyoxal | 14,5 | 23,3 | 55 | 2,156164384 | 13,20 |
| 010 K18 | male | Methylglyoxal | 14,4 | 24,8 | 61 | 4,306849315 | 14,85 |
| 013 K18 | male | Glucose | 14,1 | 24,6 | 59 | 4,304109589 | 17,00 |
| 019 K18 | female | Methylglyoxal | 14,3 | 23,8 | 57 | 4,304109589 | 13,35 |
| 020 K18 | female | Glucose | 13,9 | 23 | 59 | 4,306849315 | 16,35 |
| 022 K18 | female | Methylglyoxal | 14,8 | 23,1 | 62 | 4,304109589 | 13,05 |
| 029 K18 | female | Control | 14,3 | 23,4 | 56,5 | 4,304109589 | 14,80 |
| 035 K18 | female | Glucose | 14,4 | 23,8 | 57,5 | 4,304109589 | 12,95 |

| | | | | | | | |
|---|---|---|---|---|---|---|---|
| 044 K18 | female | Glucose | 13,7 | 23,4 | 56 | 4,298630137 | 14,05 |
| 045 K18 | female | Methylglyoxal | 14,6 | 24,2 | 58 | 4,298630137 | 13,60 |
| 047 K18 | male | Control | 13,4 | 23,5 | 62 | 4,298630137 | 15,65 |
| 049 K18 | male | Methylglyoxal | 14,6 | 24,5 | 59 | 4,295890411 | 14,25 |
| 053 K18 | female | Methylglyoxal | 14,4 | 24,5 | 62 | 4,295890411 | 15,00 |
| 054 K18 | female | Methylglyoxal | 13,9 | 24 | 62 | 4,295890411 | 15,05 |
| 059 K18 | male | Glucose | 13,5 | 24,2 | 60 | 4,293150685 | 12,65 |
| 061 K18 | male | Glucose | 13,4 | 23 | 61 | 4,290410959 | 14,40 |
| 071 K18 | male | Methylglyoxal | 13,8 | 23 | 59 | 4,287671233 | 15,25 |
| 073 K18 | female | Control | 13,9 | 22,7 | 59 | 4,282191781 | 11,85 |
| 076 K18 | female | Control | 14,6 | 23,6 | 61 | 4,24109589 | 14,20 |
| 079 K18 | male | Methylglyoxal | 13,9 | 24,9 | 59 | 4,235616438 | 13,95 |
| 088 K18 | female | Control | 13,7 | 23 | 57 | 4,210958904 | 14,30 |
| 089 K18 | female | Methylglyoxal | 14,4 | 23,3 | 57,5 | 4,208219178 | 14,35 |
| 099 K18 | female | Control | 14,5 | 24,4 | 57 | 4,194520548 | 14,20 |
| 10 16D | female | Control | 15,3 | 22,8 | 59 | 6,391780822 | 13,25 |
| 104 16D | male | Control | 15,5 | 24,1 | 61 | 6,260273973 | 15,35 |
| 104 S18 | female | Control | 14,7 | 23,5 | 58 | 4,197260274 | 12,80 |
| 106 16D | male | Control | 14,4 | 23,8 | 57 | 6,254794521 | 15,65 |
| 107 S18 | female | Methylglyoxal | 14,7 | 21,8 | 58 | 4,2 | 13,05 |
| 109 S18 | male | Methylglyoxal | 13,7 | 23,7 | 57 | 4,194520548 | 12,80 |
| 11 16D | male | Methylglyoxal | 13,9 | 24 | 59 | 6,391780822 | 15,10 |
| 112 K18 | male | Glucose | 15,1 | 23,5 | 59 | 4,194520548 | 14,40 |
| 115 S18 | female | Control | 13,4 | 23 | 61 | 4,191780822 | 13,45 |
| 119 16D | male | Glucose | 14 | 23,9 | 60 | 6,22739726 | 13,20 |
| 119 K19 | female | Methylglyoxal | 14,7 | 23,3 | 59 | 3,309589041 | 13,85 |
| 119 S18 | female | Glucose | 15,9 | 23 | 59 | 4,189041096 | 14,10 |
| 12 16D | male | Control | 13,9 | 23,7 | 60 | 6,391780822 | 15,25 |
| 121 K18 | male | Control | 14,8 | 25,9 | | 4,194520548 | 12,35 |
| 127 K18 | female | Glucose | 14,2 | 23,8 | 58 | 4,191780822 | 13,25 |
| 129 K18 | female | Glucose | 14,7 | 24,2 | 61 | 4,191780822 | 15,40 |

| | | | | | | | |
|---|---|---|---|---|---|---|---|
| 132 S18 | female | Methylglyoxal | 14,5 | 22,9 | 60 | 4,156164384 | 13,30 |
| 135 K18 | male | Methylglyoxal | 14,5 | 23 | 60 | 4,183561644 | 13,60 |
| 135 K19 | female | Glucose | 14,6 | 23,1 | 55 | 3,306849315 | 14,90 |
| 137 K18 | female | Methylglyoxal | 13,4 | 22,8 | 55 | 4,183561644 | 13,70 |
| 145 K18 | male | Methylglyoxal | 14,7 | 23,3 | | 4,145205479 | 13,05 |
| 145 S18 | female | Control | 14,4 | 23,5 | 57 | 4,139726027 | 13,75 |
| 148 S18 | male | Glucose | 15,4 | 22,7 | 58 | 4,131506849 | 10,60 |
| 151 K18 | male | Control | 14,6 | 24,1 | 57 | 4,131506849 | 14,35 |
| 153 K18 | female | Glucose | 15,7 | 23,8 | 58 | 4,134246575 | 13,65 |
| 158 K18 | female | Glucose | 13,3 | 22,3 | 57 | 4,101369863 | 11,45 |
| 173 16D | female | Methylglyoxal | 13,9 | 23,5 | 58 | 5,991780822 | 14,40 |
| 173 K19 | male | Glucose | 13,5 | 23,5 | 57 | 3,298630137 | 13,55 |
| 175 16D | male | Control | 14,3 | 24,4 | 62 | 5,994520548 | 14,30 |
| 194 16D | male | Control | 14,5 | 23,9 | 61 | 5,947945205 | 13,80 |
| 196 16D | male | Control | 14,3 | 23,9 | 60 | 5,942465753 | 14,45 |
| 205 16D | male | Glucose | 14,1 | 23,7 | 58 | 5,926027397 | 13,35 |
| 220 16D | male | Methylglyoxal | 14,9 | 24,1 | 58,5 | 5,882191781 | 12,50 |
| 222 16D | male | Control | 14,8 | 24,2 | 61 | 5,882191781 | 12,70 |
| 230 16D | female | Glucose | 14,5 | 23,6 | 60 | 5,873972603 | 12,95 |
| 234 16D | male | Glucose | 13,8 | 24,1 | 60 | 5,849315068 | 14,53 |
| 235 16D | female | Control | 14,5 | 23,4 | 58 | 5,846575342 | 14,20 |
| 242 16D | female | Glucose | 14,3 | 23,9 | 60 | 5,81369863 | 15,40 |
| 254 16D | male | Glucose | 15,3 | 24,3 | 60 | 5,769863014 | 17,25 |
| 27 16D | female | Glucose | 14,7 | 23,6 | 59 | 6,38630137 | 15,25 |
| 29 16D | male | Control | 15,1 | 24,2 | 58 | 6,383561644 | 15,45 |
| 295 K19 | male | Glucose | 14,4 | 23,8 | 57 | 3,180821918 | 12,65 |
| 37 16D | female | Methylglyoxal | 14,7 | 23,9 | 58 | 6,378082192 | 14,80 |
| 4 16D | female | Control | 14,4 | 23,8 | 60 | 6,4 | 15,00 |
| 70 16D | male | Methylglyoxal | 14,8 | 23,8 | 58 | 6,290410959 | 15,30 |
| 71 16D | female | Glucose | 14,8 | 24 | 59 | 6,287671233 | 15,75 |
| 78 16D | male | Methylglyoxal | 14,5 | 23,6 | 59 | 6,279452055 | 15,10 |

| | | | | | | | |
|---|---|---|---|---|---|---|---|
| 89 17S | female | Methylglyoxal | 14,3 | 23,5 | 62 | 5,142465753 | 14,35 |
| S520F/008 | female | Control | 17 | 26,5 | 65 | | 22,30 |

BM=body mass in grams as measured in the baseline

Tarsus, Head_beak and wing correspond to body measurements in cm.

Age_years= age in years of the birds the day of start of the experiment (see **Material and methods**). The age was not known for 4 individuals.
