## Supplementary material for "Contrasting effects of glucose and methylglyoxal supplementation on blood oxidative status, blood cells’ telomere dynamics and apoptosis in birds": ESM2 - Extra results

### Box 1. Models and their corresponding final equations.

#### Main models

**1. OXY**
lmer(OXY~Group*Month+(1|Bird_ID))
lmer(OXY~Group*Age_L*Sex+(1|Bird_ID))

**2. d-ROMs**
lmer(logROM~Group*Month*C_OXY+Month*Sex+(1|Bird_ID))
lmer(logROM~Group*C_OXY+C_OXY*Sex+Age_L*Sex+Age_L*C_OXY+Group*Age_L+(1|Bird_ID))

**3. Protein carbonyl**
lmer(logPC~Group*Month+C_OXY+Sex+(1|Bird_ID))
lmer(logPC~Group*Age_L+C_OXY+(1|Bird_ID))

**4. DNA damage**
lmer(logDNA_damage~Group*Month+(1|Bird_ID))
lmer(logDNA_damage~Group*Age_L+Sex*Age_L+(1|Bird_ID))

**5. Telomeres**
lmer(Telomeres~Group*Month+C_logDNA+C_Age_H+(1|Bird_ID))
lmer(Telomeres~Group*Age_L+C_logDNA+Age_L*C_Age_H+(1|Bird_ID))
lmer(Telomeres~C_logDNA+C_DNA_damage+(1|Bird_ID)

**6. Apoptosis**
glmmTMB(cbind(Early_apoptosis, Living_cells) ~ Group*Month*Time*Sex+C_Age_H +(1| Bird_ID/Month), family= binomial(link="logit"))
glmmTMB(cbind(Early_apoptosis, Living_cells) ~ Group*Time*Age_L+C_Age_H*Age_L + (1| Bird_ID/Month), family= betabinomial(link="logit"))

**7. Mitochondrial superoxide production**
**A. Living cells**
lmer(log10(Mean_mitosox_Cell_type) ~ Group*Month+Month*Time +Month*Time*C_N_cells-(Month:Time:C_N_cells)+ (1| Bird_ID/Month))
lmer(log10(Mean_mitosox_Cell_type) ~ Group*Age_L + Month*Time*C_N_cells-(Month:Time:C_N_cells)+ (1| Bird_ID/Month))

**B. Early apoptotic**
lmer(log10(Mean_mitosox_Cell_type) ~ Group*Month+Month*Time+ Month*C_N_cells + (1| Bird_ID/Month))
lmer(log10(Mean_mitosox_Cell_type) ~ Group*C_Age_L+Group*C_Age_L2+ Month*Time+Month*C_N_cells + (1| Bird_ID/Month))

**C. Dynamics**
lmer(Dif_Res ~ Group*Month + (1| Bird_ID))
lmer(Dif_Res ~ Group*Age_L + (1| Bird_ID))

lmer(Dif_Res ~ Group*Month + (1| Bird_ID))
lmer(Dif_Res ~ Group*Age_L + (1| Bird_ID))

#### Mortality hazard models

coxph(Surv(tstartOS, tstopROM, endpointROM)~Treatment+C_BM+C_Age_years+C_ROM)
coxph(Surv(tstartPC, tstopPC, endpointPC)~Treatment+C_BM+C_Age_years*C_PC)
coxph(Surv(tstartOS, tstopDNA, endpointDNA)~Treatment+C_BM+C_Age_years+C_DNA)
coxph(Surv(tstartTel, tstopTel, endpointTel)~Treatment+C_BM+Age+Sex+C_Telomeres+C_logDNA_amount)

### Box 2. Secondary models and their corresponding final equations.

**DNA damage on telomeres**

lmer(Telomeres~C_logDNA+C_DNA_damage+(1|Bird_ID))
lmer(Dynamics~C_DNA_damage+Month_diff+C_Age_years+C_TelbyDNA_prev+(1|Bird_ID)

**DNA damage and telomeres on apoptosis**

glmmTMB(cbind(Early_apoptosis, Living_cells) ~ Group*Month*Time*Sex+ C_DNA_damage +(1| Bird_ID/Month), family= binomial(link="logit"))
glmmTMB(cbind(Early_apoptosis, Living_cells) ~ Group*Month*Time*Sex+ C_Telomeres +(1| Bird_ID/Month), family= binomial(link="logit"))

**Whole blood glucose on oxidative status**

lmer(logROM~Month*Group+C_logGlucose*Sex+C_OXY+Month:Sex + (1| Bird_ID))
lmer(logPC~Month+Sex+C_logGlucose+C_OXY + (1| Bird_ID))
lmer(logDNA~Month*Group+C_logGlucose + (1| Bird_ID))

**Plasma glucose on oxidative status**

lmer(logROM~Month+Group+C_logPGlu*Month*Sex+C_OXY + (1| Bird_ID))
lmer(logPC~Month+C_OXY+Sex*C_logPGlu + (1| Bird_ID))
lmer(logDNA~Month*Group+C_logPGlu + (1| Bird_ID))

**Oxidative status on glycation**

lmer(Glycation~C_logPGlu+C_OXY+C_logROM+C_Age_years + (1| Bird_ID))
lmer(Glycation~+Month*C_logPGlu+C_logPC+C_OXY + (1| Bird_ID))
lmer(Glycation~Group+C_logPGlu*Month+C_logDNA+C_OXY+(1| Bird_ID))

**Oxidative status on glyoxal**

lmer(logGlyoxal~Group*Sex+Sex*C_logROM+C_Age_years+ (1| Bird_ID)
lmer(logGlyoxal~Group*Sex+C_logPC+C_Age_years+ (1| Bird_ID))

**Oxidative status on methylglyoxal**

lmer(logMG~Month+C_logROM+C_OXY+ (1| Bird_ID))
lmer(logMG~Month+C_logPC+ (1| Bird_ID))

**Oxidative status on CML**

lmer(logCML~Month+C_logROM+C_OXY+C_Age_years+ (1| Bird_ID))
lmer(logCML~Month+Sex+C_logPC+C_Age_years + (1| Bird_ID))
lmer(logCML~Month+C_logDNA*Month+C_logGlyoxal+C_OXY +C_Age_years+ (1| Bird_ID))

**Oxidative status on CEL**

lmer(logCEL~C_logROM+C_logGlyoxal + (1| Bird_ID))
lmer(logCEL~C_logPC+C_logGlyoxal + (1| Bird_ID))
lmer(logCEL~Sex*C_logDNA+C_logGlyoxal + (1| Bird_ID))

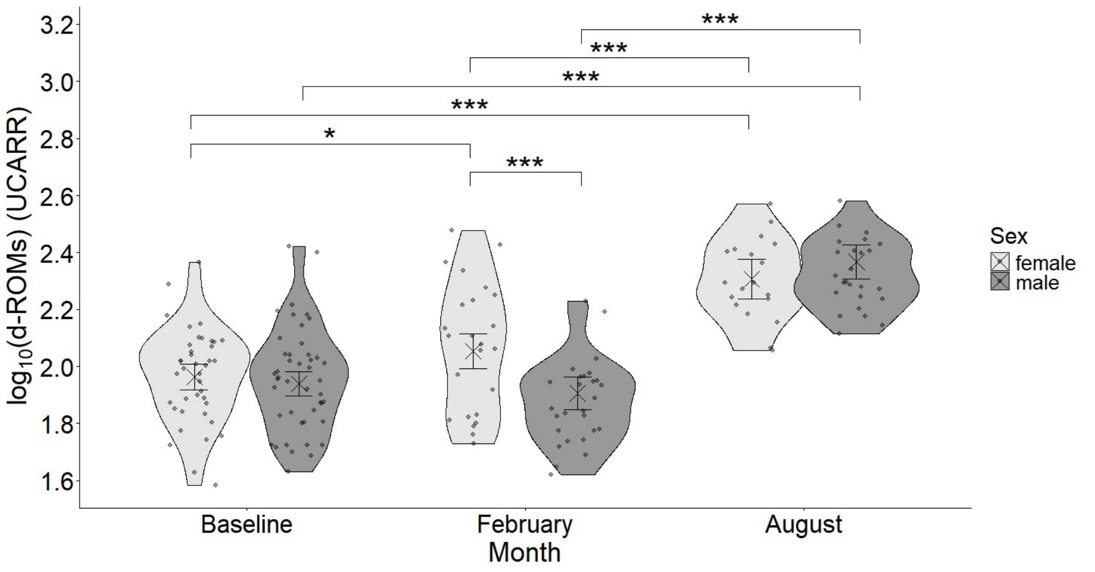


**Figure ESM2.1.** Plasma $log_{10}$ transformed d-ROMs levels in UCARR acorss months and sexes. Crosses and error bars represent model-estimated marginal means ±95% CI. Significance annotations are based on pairwise contrasts performed separately within months and within sexes.

**Table ESM2.1.** Within treatments across months marginal means contrast for total plasma organic hydroperoxid (d-ROMs in UCARR) values.

| **Group = Control:** | | | | | |
|---|---|---|---|---|---|
| **Contrast** | **Estimate** | **SE** | **df** | **t ratio** | **P value** |
| Baseline - February | -0.1098 | 0.0434 | 145 | -2.533 | 0.0329 |
| Baseline - August | -0.4220 | 0.0440 | 149 | -9.593 | <.0001 |
| February - August | -0.3122 | 0.0480 | 137 | -6.508 | <.0001 |
| **Group = Glucose:** | | | | | |
| **Contrast** | **Estimate** | **SE** | **df** | **t ratio** | **P value** |
| Baseline - February | -0.0230 | 0.0500 | 166 | -0.459 | 0.8903 |
| Baseline - August | -0.3432 | 0.0545 | 174 | -6.294 | <.0001 |
| February - August | -0.3202 | 0.0615 | 125 | -5.207 | <.0001 |
| **Group = Methylglyoxal:** | | | | | |
| **Contrast** | **Estimate** | **SE** | **df** | **t ratio** | **P value** |
| Baseline - February | 0.0476 | 0.0408 | 140 | 1.167 | 0.4749 |
| Baseline - August | -0.3924 | 0.0468 | 161 | -8.378 | <.0001 |
| February - August | -0.4400 | 0.0510 | 161 | -8.636 | <.0001 |

**Table ESM2.2.** Within sexes across months marginal means contrast for total plasma organic hydroperoxid (d-ROMs in UCARR) values.

| **Sex = female:** | | | | | |
|---|---|---|---|---|---|
| **Contrast** | **Estimate** | **SE** | **df** | **t ratio** | **P value** |
| Baseline - February | -0.0902 | 0.0371 | 157 | -2.429 | 0.0428 |
| Baseline - August | -0.3433 | 0.0413 | 171 | -8.321 | <.0001 |
| February - August | -0.2532 | 0.0453 | 142 | -5.583 | <.0001 |
| **Sex = male:** | | | | | |
| **Contrast** | **Estimate** | **SE** | **df** | **t ratio** | **P value** |
| Baseline - February | 0.0333 | 0.0351 | 146 | 0.948 | 0.6107 |
| Baseline - August | -0.4285 | 0.0363 | 151 | -11.816 | <.0001 |
| February - August | -0.4618 | 0.0406 | 132 | -11.380 | <.0001 |

**Table ESM2.2.** Within treatment across months marginal means contrast for plasma protein carbonyl levels (nmoles/mg of protein).

| **Group = Control** | | | | | |
|---|---|---|---|---|---|
| **Contrast** | **Estimate** | **SE** | **df** | **t ratio** | **P value** |
| Baseline - February | -0.02202 | 0.0498 | 116.9 | -0.442 | 0.8980 |
| Baseline - August | 0.26267 | 0.0505 | 129.1 | 5.198 | <.0001 |
| February - August | 0.28468 | 0.0536 | 110.2 | 5.313 | <.0001 |
| **Group = Glucose** | | | | | |
| **Contrast** | **Estimate** | **SE** | **df** | **t ratio** | **P value** |
| Baseline - February | -0.00832 | 0.0579 | 127.4 | -0.144 | 0.9887 |
| Baseline - August | 0.29923 | 0.0607 | 130.0 | 4.926 | <.0001 |
| February - August | 0.30755 | 0.0673 | 91.9 | 4.569 | <.0001 |
| **Group = Methylglyoxal** | | | | | |
| **Contrast** | **Estimate** | **SE** | **df** | **t ratio** | **P value** |
| Baseline - February | 0.03411 | 0.0421 | 104.8 | 0.810 | 0.6980 |
| Baseline - August | 0.25303 | 0.0445 | 111.4 | 5.691 | <.0001 |
| February - August | 0.21892 | 0.0465 | 102.9 | 4.706 | <.0001 |

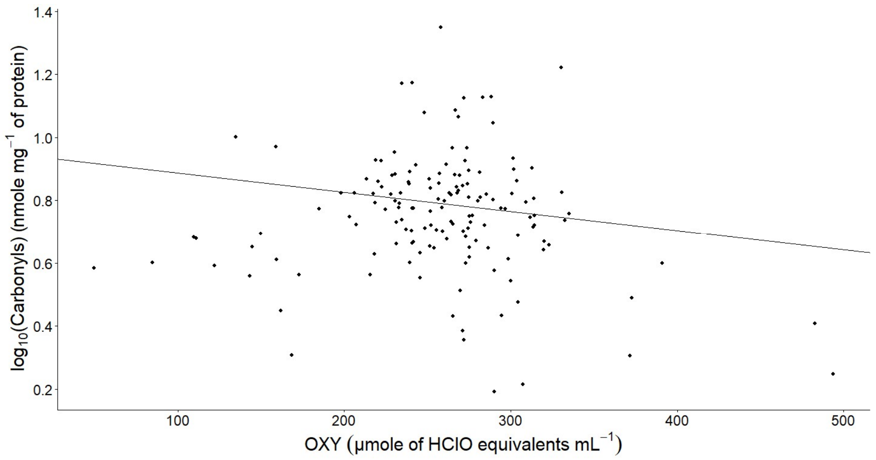


**Figure ESM2.2.** Negative effects of OXY on protein carbonyl levels.

**Table ESM2.3.** Within treatment across months contrasts of apoptosis probabilities.

| **Group = Control, Time = T0:** | | | | | | |
|---|---|---|---|---|---|---|
| **Contrast** | **Odds ratio** | **SE** | **df** | **null** | **Z ratio** | **P value** |
| November / February | 13.302 | 1.860 | Inf | 1 | 18.520 | <0.0001 |
| November / May | 44.227 | 6.500 | Inf | 1 | 25.773 | <0.0001 |
| November / August | 26.304 | 3.950 | Inf | 1 | 21.783 | <0.0001 |
| February / May | 3.325 | 0.493 | Inf | 1 | 8.101 | <0.0001 |
| February / August | 1.977 | 0.299 | Inf | 1 | 4.516 | <0.0001 |
| May / August | 0.595 | 0.093 | Inf | 1 | -3.323 | 0.0049 |
| **Group = Glucose, Time = T0:** | | | | | | |

| **Contrast** | **Odds ratio** | **SE** | **df** | **null** | **Z ratio** | **P value** |
|---|---|---|---|---|---|---|
| November / February | 3.356 | 0.639 | Inf | 1 | 6.356 | <0.0001 |
| November / May | 21.797 | 4.540 | Inf | 1 | 14.808 | <0.0001 |
| November / August | 14.998 | 3.110 | Inf | 1 | 13.040 | <0.0001 |
| February / May | 6.494 | 1.350 | Inf | 1 | 8.984 | <0.0001 |
| February / August | 4.469 | 0.928 | Inf | 1 | 7.206 | <0.0001 |
| May / August | 0.688 | 0.154 | Inf | 1 | -1.672 | 0.3386 |
| **Group = Methylglyoxal, Time = T0:** | | | | | | |
| **Contrast** | **Odds ratio** | **SE** | **df** | **null** | **Z ratio** | **P value** |
| November / February | 2.722 | 0.355 | Inf | 1 | 7.678 | <0.0001 |
| November / May | 5.233 | 0.715 | Inf | 1 | 12.105 | <0.0001 |
| November / August | 4.790 | 0.669 | Inf | 1 | 11.215 | <0.0001 |
| February / May | 1.923 | 0.262 | Inf | 1 | 4.796 | <0.0001 |
| February / August | 1.760 | 0.245 | Inf | 1 | 4.064 | 0.0003 |
| May / August | 0.915 | 0.132 | Inf | 1 | -0.613 | 0.9281 |
| **Group = Control, Time = T24:** | | | | | | |
| **Contrast** | **Odds ratio** | **SE** | **df** | **null** | **Z ratio** | **P value** |
| November / February | 8.557 | 1.200 | Inf | 1 | 15.369 | <0.0001 |
| November / May | 21.828 | 3.200 | Inf | 1 | 21.030 | <0.0001 |
| November / August | 22.689 | 3.400 | Inf | 1 | 20.815 | <0.0001 |
| February / May | 2.551 | 0.377 | Inf | 1 | 6.335 | <0.0001 |
| February / August | 2.652 | 0.400 | Inf | 1 | 6.467 | <0.0001 |
| May / August | 1.039 | 0.162 | Inf | 1 | 0.248 | 0.9946 |
| **Group = Glucose, Time = T24:** | | | | | | |
| **Contrast** | **Odds ratio** | **SE** | **df** | **null** | **Z ratio** | **P value** |
| November / February | 4.578 | 0.872 | Inf | 1 | 7.985 | <0.0001 |
| November / May | 20.040 | 4.170 | Inf | 1 | 14.417 | <0.0001 |
| November / August | 18.649 | 3.870 | Inf | 1 | 14.087 | <0.0001 |
| February / May | 4.378 | 0.911 | Inf | 1 | 7.097 | <0.0001 |
| February / August | 4.074 | 0.847 | Inf | 1 | 6.760 | <0.0001 |
| May / August | 0.931 | 0.208 | Inf | 1 | -0.322 | 0.9885 |
| **Group = Methylglyoxal, Time = T24:** | | | | | | |
| **Contrast** | **Odds ratio** | **SE** | **df** | **null** | **Z ratio** | **P value** |
| November / February | 3.253 | 0.424 | Inf | 1 | 9.045 | <0.0001 |
| November / May | 6.271 | 0.857 | Inf | 1 | 13.440 | <0.0001 |
| November / August | 6.713 | 0.938 | Inf | 1 | 13.629 | <0.0001 |
| February / May | 1.928 | 0.263 | Inf | 1 | 4.820 | <0.0001 |
| February / August | 2.064 | 0.287 | Inf | 1 | 5.208 | <0.0001 |
| May / August | 1.071 | 0.154 | Inf | 1 | 0.473 | 0.9650 |

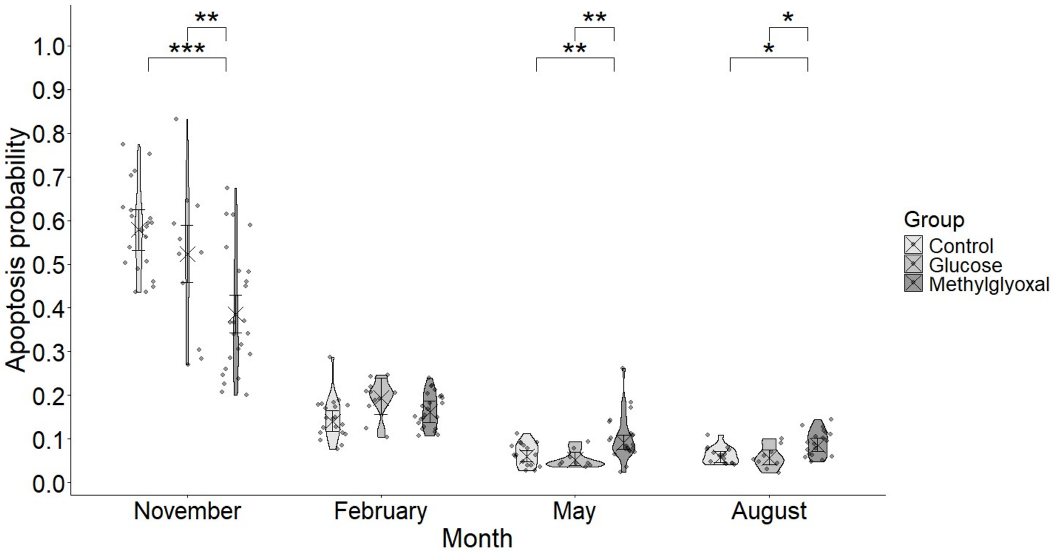


**Figure ESM2.3.** Apoptosis levels represented as a proportion of early apoptotic versus living cells along the experiment and across the treatment groups at the second measure, after 24 hours (T24). Crosses and error bars represent model-estimated probabilities ±95% CI. Significance annotations are based on pairwise contrasts performed separately within months. Comparisons within treatments across months are not included here to improve the figure visibility, due to their high number of significant results.

**Table ESM2.4.** Marginal slopes estimated for longitudinal age effects for each treatment group within each time (T0 and T24) in the apoptosis probability model.

| Group | Time | Age_L trend | SE | df | Asymptotic LCL | Asymptotic UCL |
|---|---|---|---|---|---|---|
| Control | T0 | -5.16 | 0.362 | Inf | -5.87 | -4.45 |
| Glucose | T0 | -3.91 | 0.435 | Inf | -4.76 | -3.05 |
| Methylglyoxal | T0 | -2.13 | 0.278 | Inf | -2.67 | -1.58 |
| Control | T24 | -4.53 | 0.333 | Inf | -5.18 | -3.87 |
| Glucose | T24 | -4.13 | 0.438 | Inf | -4.99 | -3.27 |
| Methylglyoxal | T24 | -2.54 | 0.279 | Inf | -3.09 | -1.99 |

**Table ESM2.5.** Across times (T0 vs T24) within treatment and months contrasts of apoptosis probabilities.

| | | | | | | |
|---|---|---|---|---|---|---|
| **Group = Control, Month = November:** | | | | | | |
| **Contrast** | **Odds ratio** | **SE** | **df** | **null** | **Z ratio** | **P value** |
| T0 / T24 | 0.834 | 0.00380 | Inf | 1 | -39.811 | <0.0001 |
| **Group = Glucose, Month = November:** | | | | | | |
| **Contrast** | **Odds ratio** | **SE** | **df** | **null** | **Z ratio** | **P value** |
| T0 / T24 | 0.823 | 0.00530 | Inf | 1 | -30.225 | <0.0001 |
| **Group = Methylglyoxal, Month = November:** | | | | | | |
| **Contrast** | **Odds ratio** | **SE** | **df** | **null** | **Z ratio** | **P value** |
| T0 / T24 | 0.818 | 0.00369 | Inf | 1 | -44.556 | <0.0001 |
| **Group = Control, Month = February:** | | | | | | |
| **Contrast** | **Odds ratio** | **SE** | **df** | **null** | **Z ratio** | **P value** |
| T0 / T24 | 0.537 | 0.00401 | Inf | 1 | -83.205 | <0.0001 |
| **Group = Glucose, Month = February:** | | | | | | |
| **Contrast** | **Odds ratio** | **SE** | **df** | **null** | **Z ratio** | **P value** |
| T0 / T24 | 1.123 | 0.00855 | Inf | 1 | 15.180 | <0.0001 |
| **Group = Methylglyoxal, Month = February:** | | | | | | |
| **Contrast** | **Odds ratio** | **SE** | **df** | **null** | **Z ratio** | **P value** |
| T0 / T24 | 0.978 | 0.00550 | Inf | 1 | -4.001 | 0.0001 |
| **Group = Control, Month = May:** | | | | | | |
| **Contrast** | **Odds ratio** | **SE** | **df** | **null** | **Z ratio** | **P value** |
| T0 / T24 | 0.412 | 0.00734 | Inf | 1 | -49.751 | <0.0001 |
| **Group = Glucose, Month = May:** | | | | | | |
| **Contrast** | **Odds ratio** | **SE** | **df** | **null** | **Z ratio** | **P value** |
| T0 / T24 | 0.757 | 0.01830 | Inf | 1 | -11.536 | <0.0001 |
| **Group = Methylglyoxal, Month = May:** | | | | | | |
| **Contrast** | **Odds ratio** | **SE** | **df** | **null** | **Z ratio** | **P value** |
| T0 / T24 | 0.980 | 0.00914 | Inf | 1 | -2.123 | 0.0337 |
| **Group = Control, Month = August:** | | | | | | |
| **Contrast** | **Odds ratio** | **SE** | **df** | **null** | **Z ratio** | **P value** |
| T0 / T24 | 0.719 | 0.01140 | Inf | 1 | -20.688 | <0.0001 |
| **Group = Glucose, Month = August:** | | | | | | |
| **Contrast** | **Odds ratio** | **SE** | **df** | **null** | **Z ratio** | **P value** |
| T0 / T24 | 1.023 | 0.02150 | Inf | 1 | 1.104 | 0.2696 |
| **Group = Methylglyoxal, Month = August:** | | | | | | |
| **Contrast** | **Odds ratio** | **SE** | **df** | **null** | **Z ratio** | **P value** |
| T0 / T24 | 1.147 | 0.01300 | Inf | 1 | 12.016 | <0.0001 |

**Table ESM2.6.** Contrasts between month marginal means estimated for MitoSox signal in living cells within each time (T0 and T24) and treatment group.

| **Group = Control, Time = T0:** | | | | | |
|---|---|---|---|---|---|
| **Contrast** | **Estimate** | **SE** | **df** | **t ratio** | **P value** |
| November - February | 1.2277 | 0.0777 | 310 | 15.800 | <0.0001 |
| November - May | 0.3600 | 0.0769 | 276 | 4.683 | <0.0001 |
| November - August | 0.7329 | 0.0765 | 255 | 9.586 | <0.0001 |
| February - May | -0.8677 | 0.0797 | 292 | -10.885 | <0.0001 |
| February - August | -0.4947 | 0.0786 | 271 | -6.292 | <0.0001 |
| May - August | 0.3729 | 0.0798 | 246 | 4.671 | <0.0001 |
| **Group = Glucose, Time = T0:** | | | | | |
| **Contrast** | **Estimate** | **SE** | **df** | **t ratio** | **P value** |
| November - February | 1.2733 | 0.0882 | 227 | 14.436 | <0.0001 |
| November - May | 0.3963 | 0.0938 | 234 | 4.225 | 0.0002 |
| November - August | 0.9376 | 0.0920 | 219 | 10.189 | <0.0001 |
| February - May | -0.8770 | 0.1000 | 222 | -8.772 | <0.0001 |
| February - August | -0.3356 | 0.0989 | 214 | -3.394 | 0.0045 |
| May - August | 0.5413 | 0.1030 | 210 | 5.267 | <0.0001 |
| **Group = Methylglyoxal, Time = T0:** | | | | | |
| **Contrast** | **Estimate** | **SE** | **df** | **t ratio** | **P value** |
| November - February | 1.1186 | 0.0663 | 237 | 16.879 | <0.0001 |
| November - May | 0.9286 | 0.0712 | 268 | 13.038 | <0.0001 |
| November - August | 0.5938 | 0.0775 | 300 | 7.662 | <0.0001 |
| February - May | -0.1900 | 0.0686 | 250 | -2.768 | 0.0307 |
| February - August | -0.5248 | 0.0745 | 280 | -7.049 | <0.0001 |
| May - August | -0.3348 | 0.0754 | 265 | -4.441 | 0.0001 |
| **Group = Control, Time = T24:** | | | | | |
| **Contrast** | **Estimate** | **SE** | **df** | **t ratio** | **P value** |
| November - February | 0.8914 | 0.0684 | 237 | 13.040 | <0.0001 |
| November - May | 0.2641 | 0.0730 | 247 | 3.616 | 0.0020 |
| November - August | 0.4442 | 0.0746 | 246 | 5.959 | <0.0001 |
| February - May | -0.6273 | 0.0752 | 253 | -8.339 | <0.0001 |
| February - August | -0.4471 | 0.0761 | 248 | -5.874 | <0.0001 |
| May - August | 0.1802 | 0.0795 | 241 | 2.265 | 0.1092 |
| **Group = Glucose, Time = T24:** | | | | | |
| **Contrast** | **Estimate** | **SE** | **df** | **t ratio** | **P value** |
| November - February | 0.9370 | 0.0979 | 297 | 9.572 | <0.0001 |
| November - May | 0.3004 | 0.0941 | 236 | 3.193 | 0.0086 |
| November - August | 0.6489 | 0.0929 | 227 | 6.985 | <0.0001 |
| February - May | -0.6366 | 0.1020 | 236 | -6.214 | <0.0001 |

| | | | | | |
|---|---|---|---|---|---|
| February - August | -0.2880 | 0.1040 | 248 | -2.761 | 0.0313 |
| May - August | 0.3486 | 0.1020 | 203 | 3.420 | 0.0042 |
| **Group = Methylglyoxal, Time = T24:** | | | | | |
| **Contrast** | **Estimate** | **SE** | **df** | **t ratio** | **P value** |
| November - February | 0.7823 | 0.0682 | 260 | 11.473 | <0.0001 |
| November - May | 0.8327 | 0.0797 | 345 | 10.446 | <0.0001 |
| November - August | 0.3051 | 0.0752 | 284 | 4.058 | 0.0004 |
| February - May | 0.0504 | 0.0834 | 365 | 0.604 | 0.9309 |
| February - August | -0.4772 | 0.0827 | 338 | -5.768 | <0.0001 |
| May - August | -0.5276 | 0.0882 | 354 | -5.984 | <0.0001 |

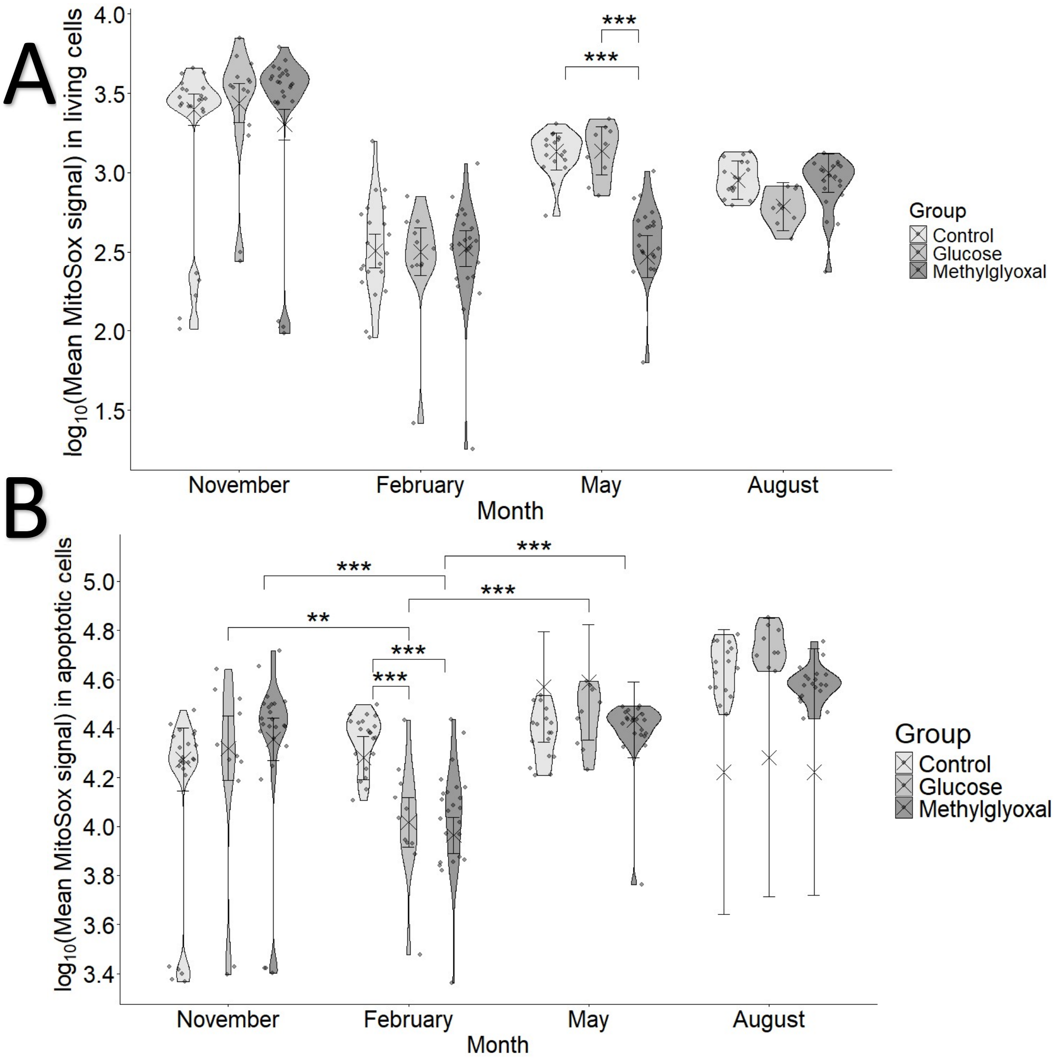


**Figure ESM2.4.** Mitochondrial superoxide production of (**A**) living cells and (**B**) early apoptotic (EA) cells, represented as the decimal logarithm of MitoSox signal along the experiment and across the treatment groups at the second measure, after 24 hours (T24). Crosses and error bars represent model-estimated marginal means ±95% CI. Significance annotations are based on pairwise contrasts performed separately within months and within treatments. As in **Figure ESM2.3**, within group (across months) comparisons are excluded from **A** in order to improve visibility.
