## Supplementary material for "Contrasting effects of glucose and methylglyoxal supplementation on blood oxidative status, blood cells’ telomere dynamics and apoptosis in birds": ESM3 - Weather

### ESM 3. Weather variations

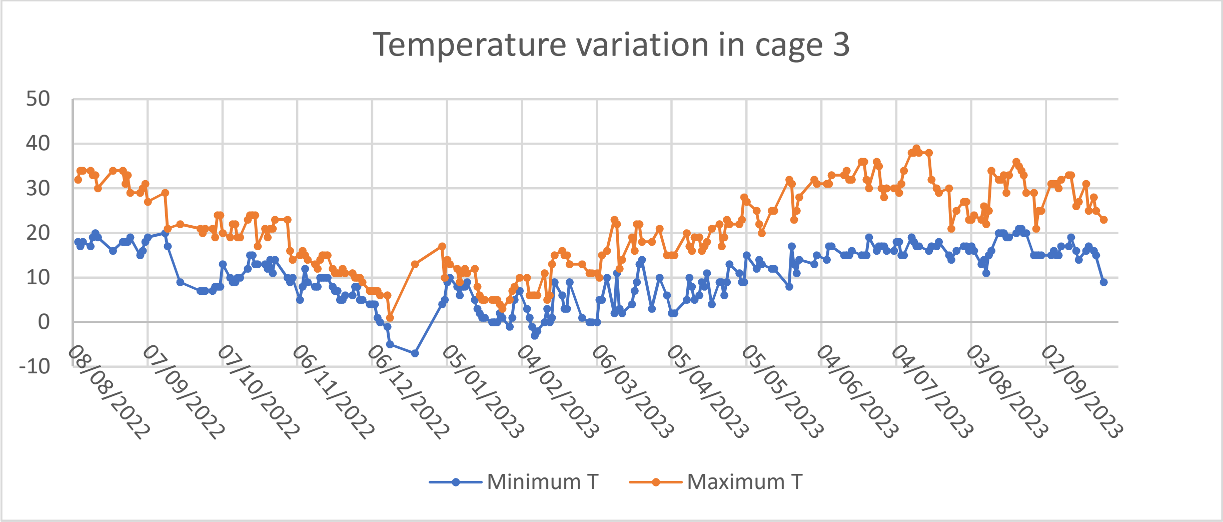


**Figure ESM3.1.** Minimum and maximum temperature variation along the year of the experiment in cage 3 (methylglyoxal supplementation group).

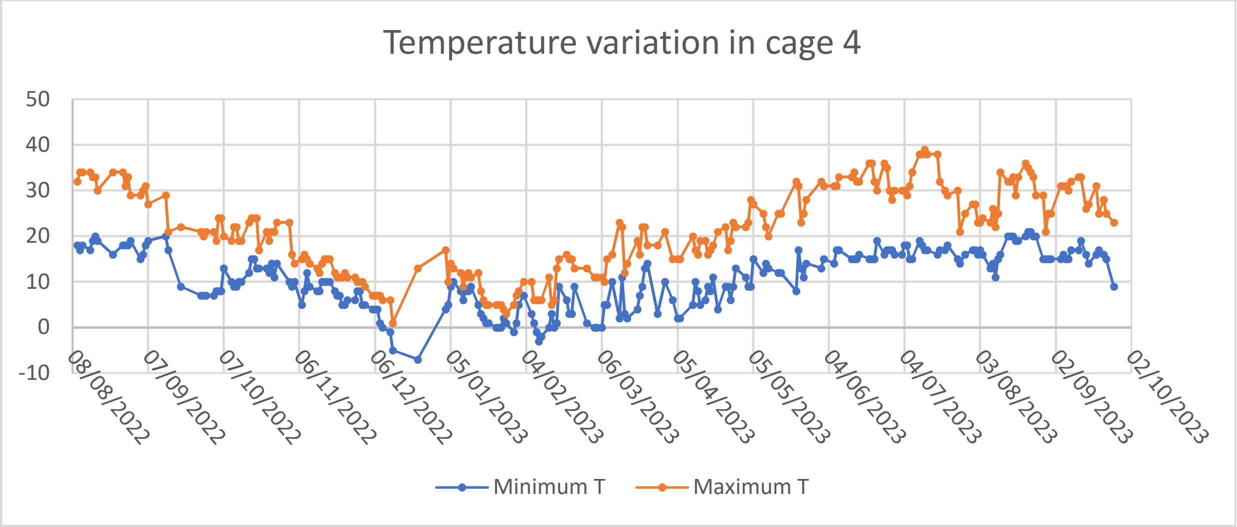


**Figure ESM3.2.** Minimum and maximum temperature variation along the year of the experiment in cage 4 (glucose supplementation group).

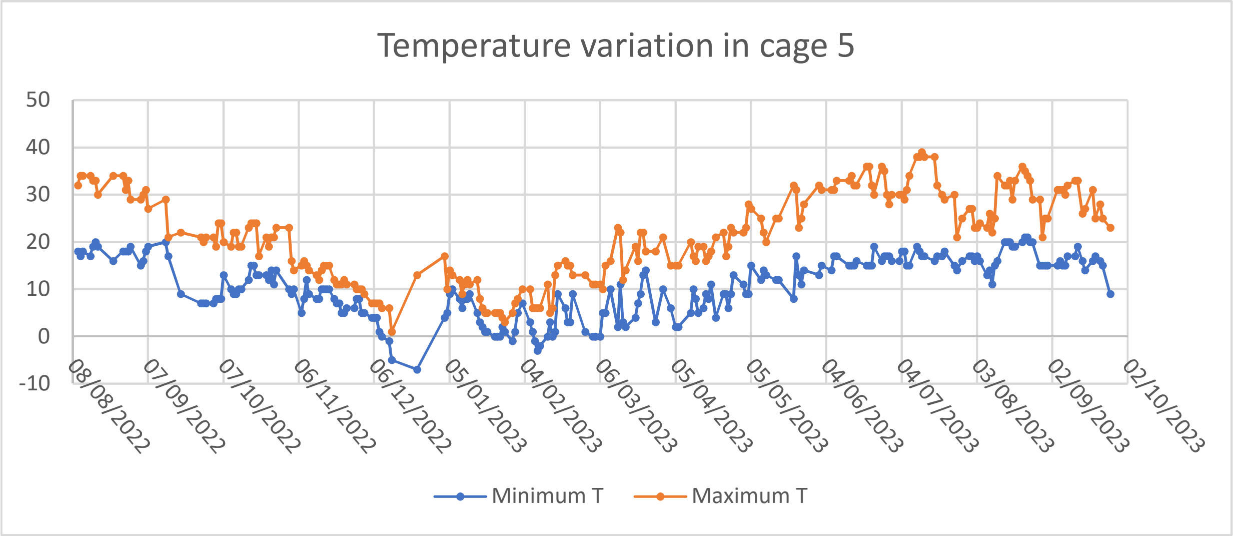


**Figure ESM3.3.** Minimum and maximum temperature variation along the year of the experiment in cage 5 (control group).
